# RegEvol: detection of directional selection in regulatory sequences through phenotypic predictions and phenotype-to-fitness functions

**DOI:** 10.1101/2025.11.26.690685

**Authors:** Alexandre Laverré, Thibault Latrille, Marc Robinson-Rechavi

## Abstract

Regulatory DNA controls when and where genes are expressed, making it a key driver of phenotypic evolution. Yet detecting selection in non-coding regions remains difficult, as most approaches rely on sequence conservation or changes in substitution rates rather than molecular effects. RegEvol bridges this gap by linking machine learning-based predictions of transcription factor binding to explicit evolutionary models. It uses the distribution of predicted mutational effects to infer fitness functions under different evolutionary scenarios including random drift, stabilising selection, and directional selection. Through maximum-likelihood estimation, it identifies the regime that best explains observed changes along a lineage from an ancestral sequence. When substitution numbers are limited, such as along short evolutionary branches, likelihood differences can be aggregated across sets of regulatory elements to increase statistical power. RegEvol corrects biases that affected previous tests based on machine learning of transcription factor binding, while remaining conservative across different levels of divergence. Applied to over 3 million Drosophila melanogaster regulatory regions, we identify 5.1% under directional selection, enriched near reproductive and immune genes. Applying the aggregation strategy to human CTCF binding across tissues reveals enrichment of directional signals in nervous and male reproductive systems. The framework is readily applicable to experimentally detected regulatory elements with alignable ancestral sequences and is adaptable to future advances in understanding regulatory function, providing a powerful basis for investigating adaptation in non-coding regions.

## Introduction

The evolution of *cis*-regulatory sequences plays a central role in shaping phenotypic diversity and adaptation across species. Regulatory evolution has been proposed as a significant factor in morphological and physiological change since King and Wilson’s well-known suggestion that many phenotypic divergences between closely related species may be caused by regulatory changes rather than differences in protein coding (King and Wilson 1975). This view is now supported by abundant evidence, with the majority of variants associated with phenotypic traits and human diseases falling within non-coding regulatory elements, such as enhancers and promoters (Maurano et al. 2012; F. Zhang and Lupski 2015; P. H. Lee et al. 2018). These sequences attract transcription factors (TFs) that bind to DNA by recognising specific sequence motifs, enabling precise gene expression patterns. This motif-based binding provides a direct link between genotype and phenotype, where the studied phenotype is a quantitative molecular trait. Moreover, mutations affecting cis-regulatory sequences that alter TF binding have been shown to contribute to macroscopic phenotypic diversity within and between species (Wittkopp and Kalay 2012; Albert and Kruglyak 2015), serving as the basis for sequence-based models of regulatory function (Lai et al. 2019; Sokolova et al. 2024).

Despite their functional importance, studying the evolutionary forces on regulatory sequences remains a major challenge. Unlike protein-coding sequences, where the ratio of nonsynonymous to synonymous substitutions (dN/dS) provides a framework to detect selection (Tanaka and Nei 1989; Yang 1998; Yang 2014), regulatory sequences lack a comparable metric. This is partly due to the absence of a universal “regulatory code” that would allow for a systematic interpretation of mutations’ functional effects in non-coding DNA (Kim and Wysocka 2023). As a result, it remains difficult to distinguish between neutral drift, stabilising selection, and adaptive evolution in regulatory regions. The majority of commonly applied approaches to detect selection in non-coding DNA are based on substitution rate variation along phylogenies. Tools such as PhastCons (Siepel et al. 2005) and PhyloP (Pollard et al. 2010) identify conserved and accelerated regions by comparing observed substitution rates to neutral expectations. These approaches have been widely used to identify deeply conserved elements that are likely to be functionally important, and can allow for branch-specific rate shifts. However, they remain limited in several significant aspects. First, substitution rate-based methods are inherently indirect: they infer selection from sequence conservation or acceleration patterns, without modelling the functional consequences of mutations. As a result, they can be confounded by non-adaptive processes such as biased gene conversion, variation in mutation rates, or demographic history (Galtier and Duret 2007). Second, because they are based on conservation across lineages, they can be poorly suited for detecting selection in fast-evolving or recently gained regulatory elements. Indeed, most cis-regulatory elements are not conserved at large evolutionary distances (Villar et al. 2015), and can maintain conserved regulatory activity despite substantial sequence divergence (Wong et al. 2020; Phan et al. 2025). Finally, they often rely on predefined neutral proxies, such as 4-fold degenerate sites or repetitive elements. These may not accurately reflect local mutational or selective backgrounds, especially in non-model species with limited annotations. Recent studies have attempted to enhance these models by incorporating phylogenetic structure and functional annotations. For example, a probabilistic framework has been developed to model the turnover of cis-regulatory elements using ChIP-seq data and phylogenetic hidden Markov models (Dukler et al. 2020). Similarly, Yan et al. introduced PhyloAcc-GT, a Bayesian method that detects substitution rate shifts in conserved noncoding elements while accounting for gene tree discordance (H. Yan et al. 2023). These approaches represent important advances in modeling substitution rate variation, but they still rely on the underlying premise that conservation of function is coupled to conservation of sequence.

To overcome these limitations, recent efforts have focused on building genotype-to-phenotype maps that directly link regulatory sequence variation to functional output at multiple molecular levels, including chromatin accessibility, transcription factor binding affinity, and downstream gene expression. Massively parallel reporter assays have enabled the systematic measurement of the effects of thousands of mutations on enhancer activity, providing empirical fitness landscapes for regulatory elements (Patwardhan et al. 2012; Smith et al. 2013; Gallego Romero and Lea 2023). These data have been used to train machine learning models that predict TF binding, chromatin accessibility, and gene expression from DNA sequence alone (Vaishnav et al. 2022; Krieger et al. 2022). These approaches demonstrate that the functional impact of mutations in regulatory DNA can be inferred directly from sequence, providing a powerful alternative to conservation-based methods. Building on this conceptual foundation, Liu and Robinson-Rechavi (2020) previously developed a machine learning-based method to detect positive selection on regulatory sequences by comparing observed changes in predicted TF binding affinity to a null distribution of random mutations. This approach leveraged gapped k-mer support vector machines (gkm-SVMs) trained on ChIP-seq data to predict the impact of substitutions on TF binding. It can infer whether a regulatory sequence has evolved under positive selection by comparing the predicted change in binding affinity (ΔSVM) to a null distribution generated by in silico mutagenesis. This method offers clear advantages over traditional rate-based approaches, as it avoids reliance on predefined neutral sites, can be applied to individual regulatory elements, and models the functional impact of mutations directly from sequence. Yet despite these strengths, several important limitations have emerged. One is ascertainment bias in ChIP-seq data, where enrichment for high-affinity binding sites can inflate false-positive rates in selection tests (Jiang and Zhang 2024). Another is that the method’s accuracy declines with divergence, since its non-parametric null model becomes unreliable over larger evolutionary distances.

In addition to these methodological issues, there is a conceptual difference between conservation-based approaches and those based on biochemical results such as ChIP-seq. Function in biology, and especially for genome regions, can be defined either by the causal role of an element or by its selected effect (Doolittle et al. 2014). Sequence conservation over long evolutionary time is predictive of selected effect, as natural selection is necessary to counteract mutation (Graur et al. 2013). Functional genomics such as ChIP-seq is predictive of causal role, since they show activity but do not demonstrate that it is selected. These are not exclusive, as regions with selected effects should have a causal role. Moreover, regions can have a selected effect yet be lowly conserved, e.g. novel genes (Keeling et al. 2019) but also regulatory sequences (Phan et al. 2025). Thus while evolutionary modeling aims to study function as selected effect, evidence for function as a causal role provides important evidence.

Here we present RegEvol, a new framework building on our previous work by combining machine learning-based predictions of mutational effects with explicit population-genetic models of selection. Rather than relying on substitution rate shifts or permutation-based tests, we directly model the fitness effects of mutations in regulatory sequences. Using a trained sequence-to-function model, our approach predicts the functional impact of all possible point mutations in ChIP-seq-defined regulatory regions. This generates genotype-to-phenotype maps that describe how sequence variation affects transcription factor binding. These predictions are then incorporated into a population-genetic framework that uses phenotype-to-fitness functions to determine the fitness effect of phenotypic changes. We evaluate the performance of this method through simulations and compare it with our previous one. Finally, we apply it to diverse empirical datasets in *Drosophila melanogaster*, and to human and mouse organ-specific TF binding, to study the evolutionary dynamics of regulatory sequences.

## Results

### 1 – SVM predictions capture functional signals beyond sequence conservation

We analysed raw public ChIP-seq datasets and trained *de novo* gapped k-mer support vector machine (gkm-SVM) models to distinguish transcription factor (TF)-bound regions identified by ChIP-seq from random genomic sequences matched for length, GC content, and repeat composition (D. Lee 2016). These models learn to recognize specific combinations of short DNA motifs (k-mers) enriched in TF-bound regions, thereby capturing the sequence context and features that define functional regulatory elements. Despite the rise of deep learning frameworks such as DeepSEA (Zhou and Troyanskaya 2015) and BPNet (Avsec et al. 2021), gkm-SVM remains among the most interpretable and TF-specific approaches, with performance comparable to deep models when trained on individual TF datasets (Tognon et al. 2025; Vorontsov et al. 2025). Moreover, gkm-SVM directly quantifies sequence-level contributions, enabling efficient in silico mutagenesis at base-pair resolution. Once trained for each dataset (specific species, tissue, and TF), the model assigns an SVM score to any DNA sequence, reflecting its predicted similarity to the ChIP-seq peak class based on k-mer composition.

We first confirmed the biological relevance of these predictions by analysing how SVM scores vary along individual ChIP-seq peaks. While previous work using this SVM approach focused on peak-level scores (Liu and Robinson-Rechavi 2020), we computed position-wise SVM scores using a sliding window approach. This allowed us to assess binding affinity prediction at the scale of individual nucleotides and to relate them to precise molecular and evolutionary metrics. A representative example is shown in Figure 1A for a sequence bound by CEBPA in *Homo sapiens* liver, which reveals two distinct regions of elevated scores, each approximately 10 bp in length, corresponding to predicted TF binding sites. These high-scoring regions matched the canonical human CEBPA motif from the JASPAR database (Castro-Mondragon et al. 2022), suggesting that the model accurately identifies functional binding sites within regulatory elements. To generalise this observation, we extracted the top 100 10-mers with the highest SVM weights from the CEBPA model and compared them to known motifs using the TomTom tool from the MEME Suite (Gupta et al. 2007; T. L. Bailey et al. 2009). The top-scoring matches corresponded to the CEBPA motif and related forkhead TFs (Supplementary Table 1), confirming that the model captures biologically meaningful sequence features associated with TF binding.

**Figure 1:**
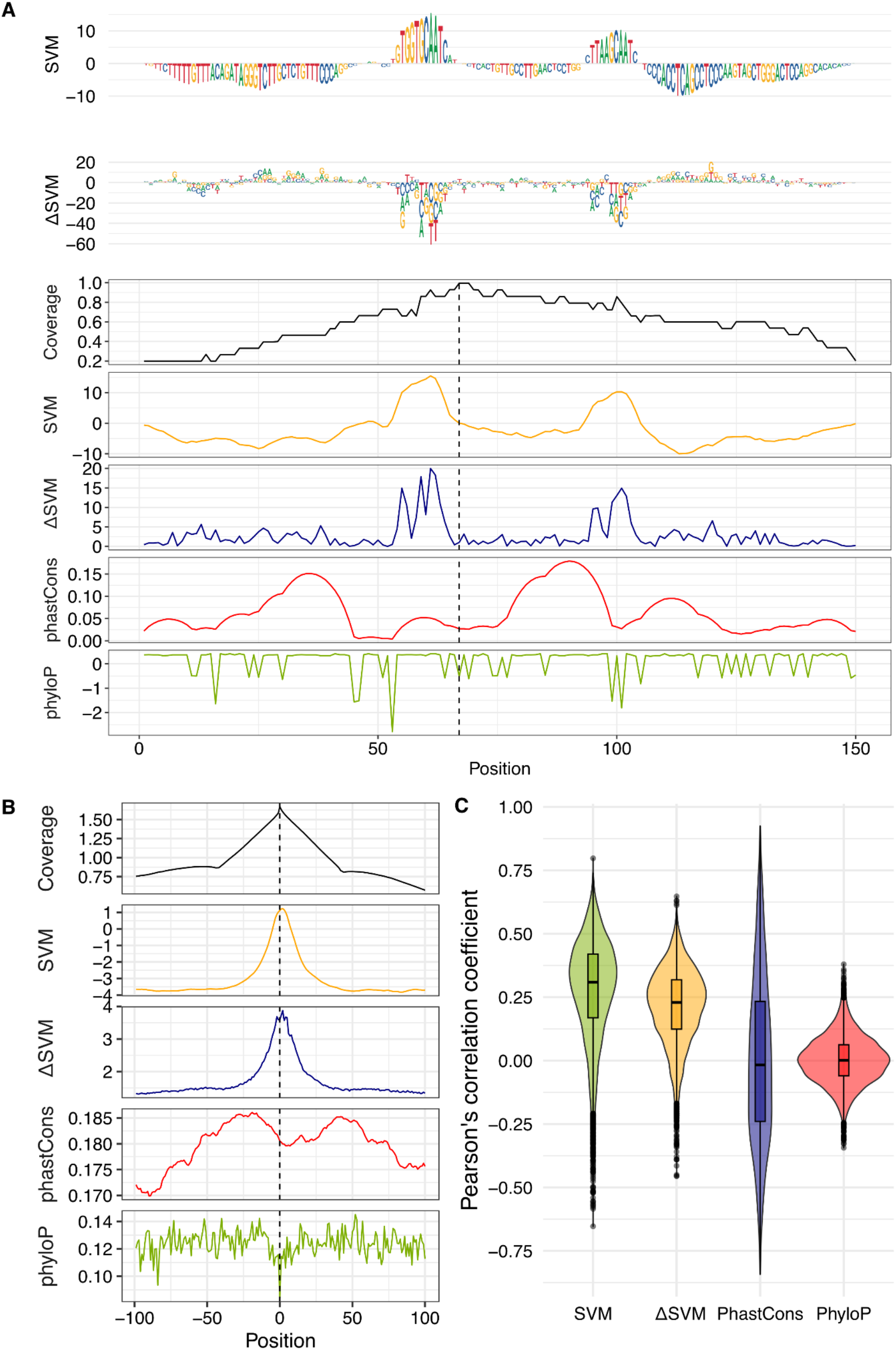
SVM-based predictions capture functional signals of transcription factor binding. **A)** SVM scores computed across a CEBPA ChIP-seq peak 188 in *Homo sapiens*. Each letter represents a nucleotide, with font size proportional to the SVM score. Two regions of elevated scores match the canonical CEBPA binding motifs (JASPAR). The summit of the peak is shown by a dotted line. **B)** Average profiles of normalised read coverage (red), SVM prediction scores (yellow), absolute ΔSVM values (green), phastCons score (blue), and phyloP score (purple), centred on the summits of CEBPA ChIP-seq peaks. **C)** Distribution of Pearson correlation coefficients between each metric (SVM, ΔSVM, phastCons, phyloP) and the base level read coverage profile across individual CEBPA peaks.

We then quantified the predicted impact of point mutations using ΔSVM, defined as the change in SVM score after a single-nucleotide substitution. At each position in the sequence, we introduced all possible point mutations in silico and computed the change in predicted binding affinity. The absolute ΔSVM values reflect the magnitude of predicted functional disruption caused by each mutation. To assess how well these metrics reflect TF binding intensity, we computed normalised read coverage for each base within CEBPA ChIP-seq peaks, using it as a proxy for bound TF abundance. All peaks were centred on their summits (the position of maximum read coverage) and aligned to produce an average signal profile. We then calculated SVM scores, ΔSVM values, and conservation metrics (phastCons and phyloP scores) at each aligned position. Both SVM and ΔSVM scores exhibit a sharp peak at the summit, closely mirroring the ChIP-seq signal (Figure 1B). This indicates that mutations in these regions are predicted to most strongly affect TF binding, either directly or indirectly. In contrast, phastCons and phyloP scores lack such enrichment and have a lower dynamic range, consistent with sequence conservation reflecting broader functional constraints rather than fine-scale, lineage-specific TF-DNA interactions. Indeed, both SVM and ΔSVM scores are positively correlated with read coverage over individual peaks (median Pearson’s correlation coefficient = 0.31 and 0.22 respectively; Fig 1C), while phastCons and phyloP scores are largely uncorrelated (median Pearson’s correlation coefficient = 1.8e-2 and 2.7e-3). This confirms that SVM-based metrics capture TF binding intensity more accurately than conservation-based scores. Notably, ΔSVM allows us to capture an evolutionary signal from the biochemical signal of ChIP-seq, thus providing a potential link between the causal role and selected effect definitions of function. These patterns were consistent across multiple TFs and species (Supplementary Figure 1). Importantly, the strength of the base-level SVM–ChIP correlation increases with peak quality, and exclusion of the lowest-quality peaks substantially improves these correlations (Supplementary Figure 2), indicating that modest or negative correlations primarily reflect technical variability rather than limitations of the modelling.

Finally, we examined the consistency of model predictions across species and TFs using mammalian liver ChIP-seq datasets (Supplementary Figure 3-4). Clustering based on k-mer scores revealed that peaks grouped primarily by TF identity rather than species, indicating that the models capture conserved binding preferences. An exception was observed for HNF4 and FOXA1, which clustered by species rather than individual TF. These two transcription factors are known to co-occupy regulatory regions in the genome, acting cooperatively and sequentially to activate liver-specific gene expression (Horisawa et al. 2020), and thus there is little sequence signal differentiating their individual binding from each other.

### 2 -Defining genotype-to-fitness maps for TF-ChIP-seq peaks

To leverage this functional prediction by SVMs and investigate the evolutionary forces acting on ChIP-seq peaks, we developed RegEvol. This framework quantifies selection on regulatory sequences by integrating predictive models of binding affinity with evolutionary theory. This approach is illustrated in Figure 2 and detailed mathematical formalism is described in Supplementary Materials.

**Figure 2.**
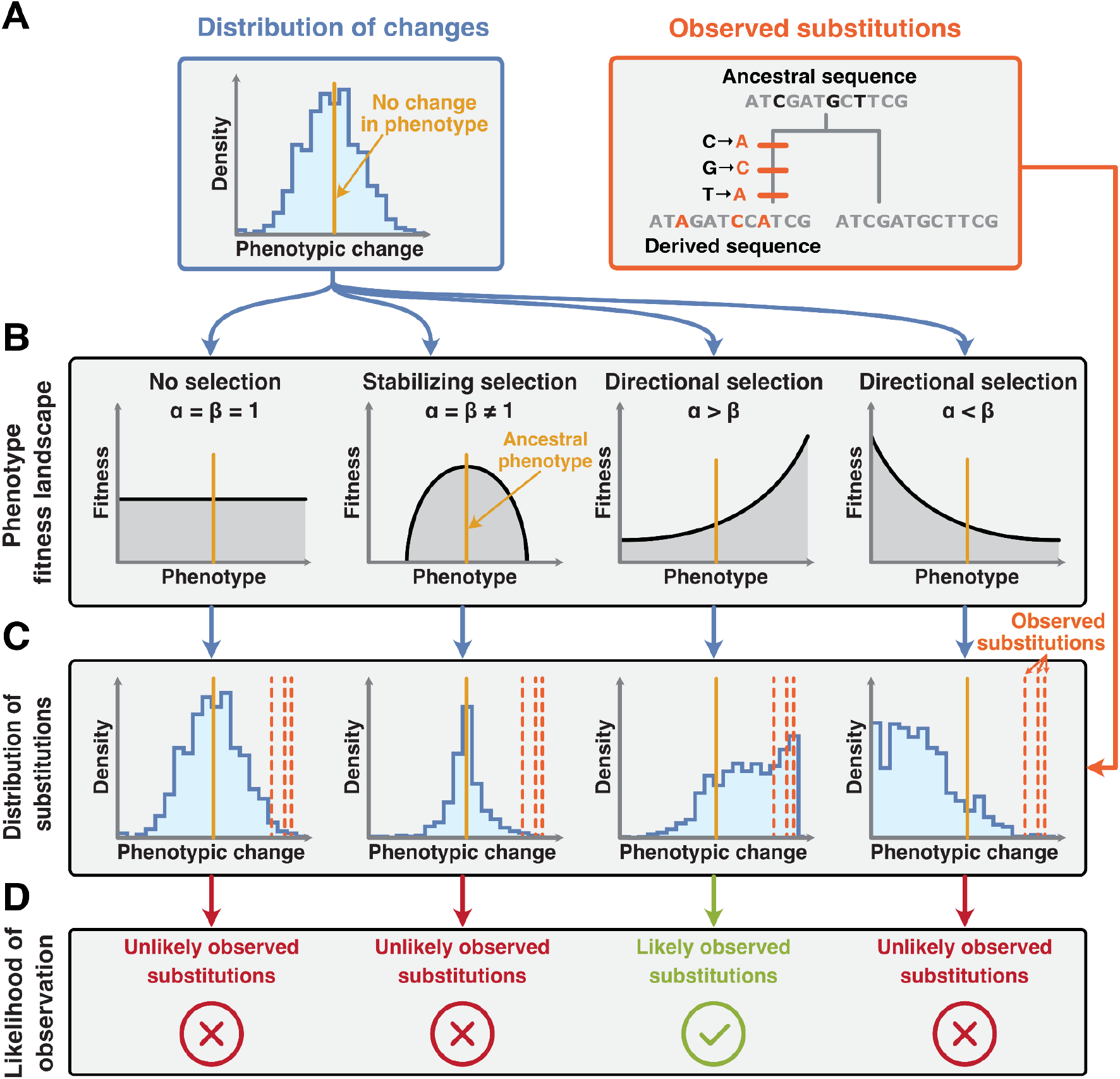
Outline of RegEvol: from genotype to fitness. **A)** Distribution of phenotypic effects (DPE) for all possible point mutations within a ChIP-seq peak (left). The observed substitutions relative to the ancestral sequence represent a subset of this distribution and are used to infer the peak’s evolutionary regime (right). B) Definition of nested selective regimes using Beta distributions parameterized by α and β, which define the shape of the underlying fitness landscape. C) Expected distributions of all possible substitutions under each phenotypic fitness landscape. Vertical orange lines indicate the observed substitutions. D) Likelihood of the observed substitutions under each selective regime. In this example, because the substitutions are biased toward positive phenotypic change, the model of directional selection for increased binding affinity is the most likely.

The first step involves constructing a genotype-to-phenotype map for each ChIP-seq peak, describing how sequence variation affects transcription factor binding affinity (i.e., molecular phenotype). Building on the base pair-resolution SVM scores introduced above, this is achieved by performing in silico mutagenesis on all possible point mutations within the peak and computing the change in predicted binding affinity for each mutation using ΔSVM. The resulting ΔSVM values define a Distribution of Phenotypic Effects (DPE), which captures the range of potential binding changes accessible from the ancestral sequence in one point mutation (Figure 2.A, left). Each DPE is specific to a given individual peak and provides a quantitative representation of its local genotype-to-phenotype landscape, where the phenotype is local TF binding. Substitutions that occurred along the focal lineage are identified by comparing the reference sequence to its inferred ancestral state, using whole-genome alignments (Figure 2.A, right). The corresponding ΔSVM is computed for each observed substitution, representing the phenotypic effect of the mutations that were actually fixed in the lineage. These observed substitutions can be viewed as a subsample of the whole mutational landscape defined by the DPE.

Three nested evolutionary scenarios are defined to model how selection may have shaped these substitutions, each corresponding to a different phenotype-to-fitness map (Figure 2.B). These maps transform the predicted phenotypic effects into fitness consequences, effectively modelling a distribution of fitness effects (DFE) specific to each regulatory element. Rather than assuming a fixed DFE, we derive it from a mechanistic model of molecular function, consistent with the population genetic framework proposed by Eyre-Walker and Keightley (2007). These models are parameterised using Beta distributions. The neutral model assumes that there is no selection on binding affinity, with mutations having a probability of fixation that does not depend on ΔSVM. This is captured by a flat fitness landscape, corresponding to a uniform Beta distribution with α = β = 1. Importantly, variation in mutation and substitution rates across the genome is accounted for in all evolutionary scenarios (Materials & Methods). The stabilising selection model assumes that the ancestral binding affinity is optimal, disfavoring mutations that deviate from this optimum. This is modelled by a symmetric Beta distribution centred at ΔSVM = 0, with α = β ≠ 1. The shape of the distribution reflects the strength of selection, with higher values of α and β indicating stronger selection. The directional selection model allows for asymmetric fitness landscapes, favouring mutations that increase or decrease binding affinity. This is modelled by an asymmetric Beta distribution with α ≠ β, allowing the fitness peak to shift away from the ancestral state. Each fitness function is combined with the DPE to generate an expected distribution of fixed mutations under the corresponding selection regime. The observed substitutions are then compared to these expectations to assess which model best explains the data.

To infer the most likely evolutionary scenario for each peak, model parameters are estimated by maximising the likelihood of the observed substitutions, given the DPE (Figure 2.C). The selection coefficient for each mutation is computed as the log-ratio of fitness between the derived and ancestral states. Fixation probabilities are calculated accordingly, and the likelihood of the observed substitutions is computed under each model. Likelihood ratio tests are then used to compare models and determine the best-fitting scenario for each peak. The model with the highest likelihood, adjusted for complexity, is selected as the most probable evolutionary explanation: neutral evolution, stabilising selection, or directional selection (Figure 2.D).

### 3 -Detection of Directional selection on simulated peaks evolution

To evaluate the performance of our likelihood-based method, we simulated the evolution of reconstructed ancestral sequences of ChIP-seq peaks under three evolutionary scenarios: random sampling (i.e., drift), skewed towards mutations with low effect (i.e., stabilising selection), and skewed towards a directional change (i.e., directional selection). For each scenario, we generated synthetic sequences by sampling substitutions according to their mutation probability, predicted phenotypic effect (ΔSVM), and fixation probability under a specified selection model. Selection strength was controlled by adjusting the parameters of the Beta distribution used to define the phenotype-to-fitness map. RegEvol produced well-calibrated p-values and controlled false discovery rates across scenario comparisons, demonstrating that likelihood-based inference accurately reflects statistical confidence (Supplementary Figure 5). Across all scenarios, the method showed high accuracy in identifying the correct evolutionary model (Figure 3.A). The method showed strong power to detect directional selection, with true positive rates (TPR) of 0.90 for directional selection for increased binding affinity and 0.77 for directional selection for decreased binding affinity, under moderate selection strength (α=10) Notably, the test was conservative: false positives (i.e cases where directional selection was inferred under neutrality or stabilising selection) remained at 1e-4 at a FDR threshold of 1%.

**Figure 3:**
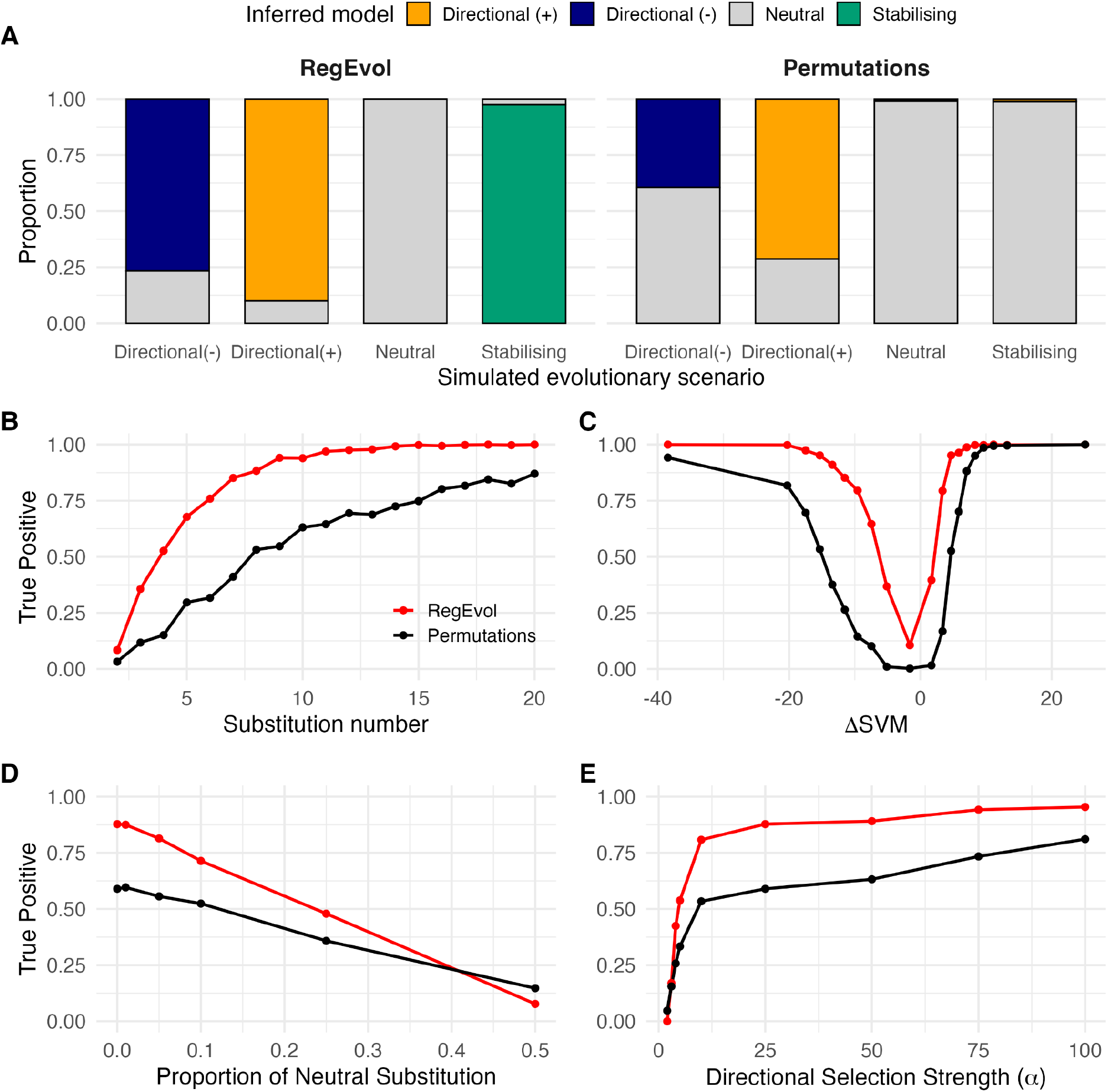
Detection of directional selection on simulated evolution of drosophila CTCF peaks. **A)** Proportion of peaks simulated under Directional, Stabilising and Random scenarios and detected by RegEvol and the Permutations Test as evolving under directional selection for positive (+) changes (orange), directional selection for negative (-) changes (blue), stabilising selection (green) and random drift (grey). **B-E)** True Positive rate of peaks simulated under directional selection (toward positive (+) or negative (-) changes) detected by RegEvol (red) or the Permutation Test (black), as a function of **B)** number of substitutions per peak, **C)** ΔSVM (quantile 5%), **D)** proportion of random substitutions, and **E)** selection strength (α parameter). *N* = 10,000 simulated peaks for each evolutionary scenario; FDR ≤ 0.01.

We quantified the factors influencing detection power by simulating the evolution of sequences under directional selection from *D. melanogaster* CTCF peaks (Figure 3.B-E), and contrasted RegEvol to our previous permutation test (Liu and Robinson-Rechavi 2020). RegEvol outperformed the permutation test across all ranges of conditions, with both higher sensitivity and specificity. As expected, detection rates increased sharply with substitution counts (Figure 3.B). Peaks with fewer than five substitutions rarely reached significance in the permutation test, showing a TPR below 0.3, and remained below 0.75 for RegEvol. The TPR for RegEvol plateaued near its maximum after about ten substitutions, whereas the permutation test increased more gradually and never reached full detection, even at 20 substitutions per peak. With only two substitutions per peak, RegEvol’s TPR remained low even under strong directional selection (TPR=0.24 at *α*=100) but reached its maximum plateau at four substitutions (Supplementary Figure 6). This reflects the conservative nature of our likelihood framework, which requires consistent phenotypic shifts across multiple substitutions to confidently reject the null model.

We observed an asymmetry in detection power for directional selection: peaks evolving toward decreased binding affinity (i.e., negative ΔSVM) were less frequently detected (Figure 3.A and C). This reflects the typical asymmetry of the DPE across species and transcription factors, with a higher proportion of mutations showing negative effects (Supplementary Figure 7; mean ΔSVM in DPE for Drosophila CTCF peaks=-0.33, sd=1.44). Moreover, as expected, detection rates were low for simulated peaks with the smallest ΔSVM (i.e., those causing little to no change in predicted binding affinity). However, while the permutation test performs poorly in this regime, RegEvol remains capable of detecting consistent directional shifts even when individual substitutions have modest effects. This is because the likelihood framework integrates directional consistency across multiple substitutions, rather than relying solely on the sum of their effects. This distinction highlights the strength of RegEvol in identifying subtle but systematic evolutionary trends.

To test the robustness of our method, we introduced a proportion of randomly fixed substitutions (i.e., mutations sampled without regard to their ΔSVM) to simulate evolutionary noise. This reflects realistic scenarios where drift is strong or multiple factors beyond the focal TF impact the evolution of regulatory elements. As expected, detection power declined with increasing proportions of such substitutions, dropping below 0.5 when one-quarter were random (Figure 3.D). RegEvol outperformed the permutation test until half of the substitutions were random, where the two methods have TPR below 0.25. A similar pattern was observed when varying directional selection strength: both methods improved with stronger selection and plateaued at comparable levels, with RegEvol maintaining superior performance (Figure 3.E).

### 4 -Specificity and Ascertainment Bias

We also evaluated the sensitivity of both methods to extreme-effect substitutions, sequence divergence, and potential ascertainment bias. The permutation test, by design, is highly responsive to large ΔSVM values. While this can be advantageous for detecting isolated, high-impact substitutions, it also makes the test more prone to false positives when such mutations are fixed by chance. In simulations where substitutions were drawn independently of their ΔSVM (Figure 4.A), the permutation test showed an overall low false positive rate (FPR=0.01) but a sharp increase for extreme ΔSVM values, particularly positive ones (quantile 1% FPR=0.15; quantile 99% FPR=0.39). A similar pattern was observed in simulations under stabilising selection, where substitutions are constrained to low ΔSVM values (Figure 4.B, quantile 99% FPR=0.26). In contrast, RegEvol remained stable and conservative across all simulations, as it integrates information across all substitutions and emphasizes consistent directional trends rather than outliers (maximum FPR across quantile=1e-4).

**Figure 4.**
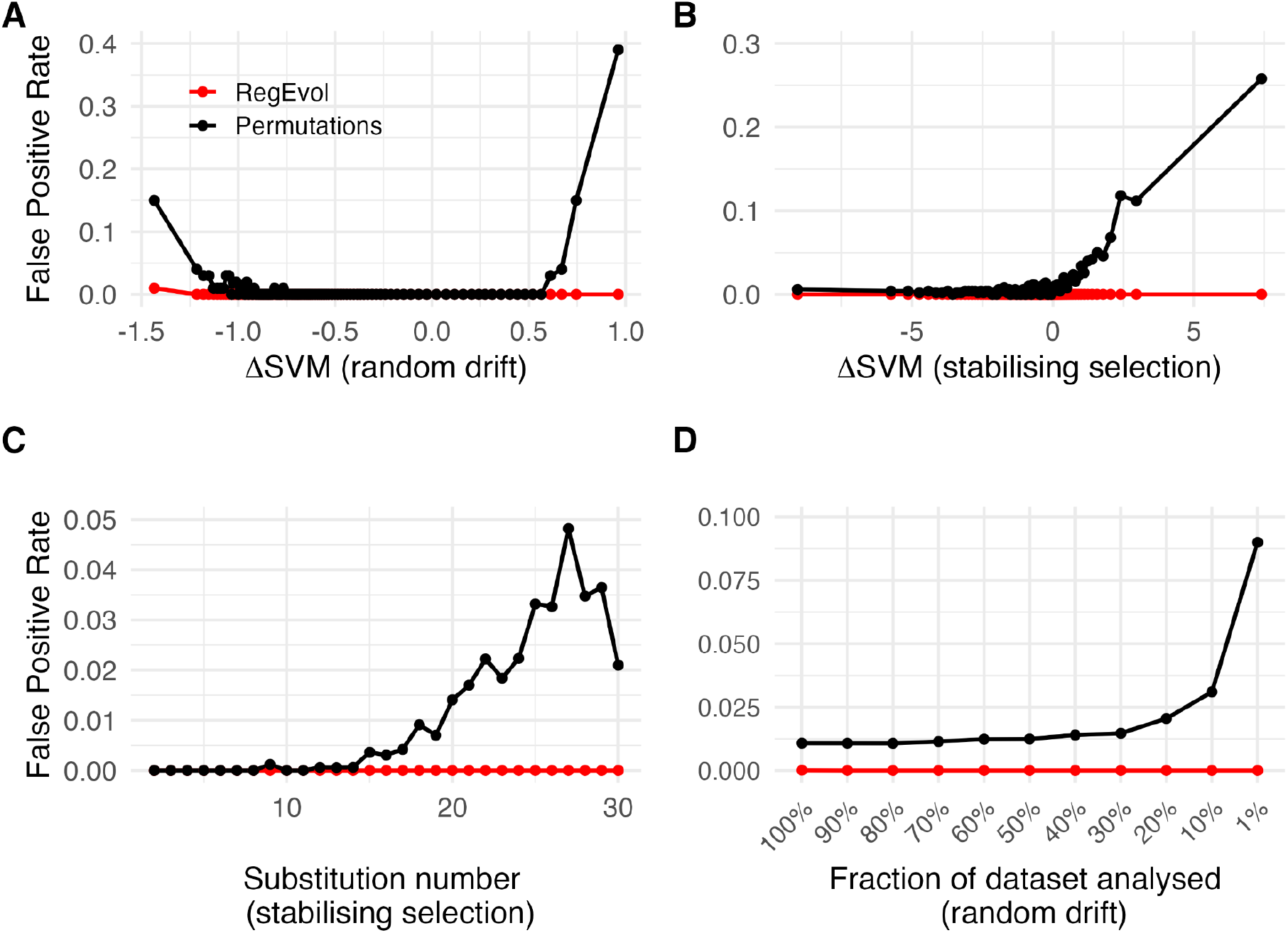
Sensitivity to extreme substitutions and robustness to ascertainment bias. Proportion of simulated *Drosophila* CTCF peaks detected under directional selection using the Permutation Test (black lines) and RegEvol (red lines). Substitutions were either drawn randomly (A, D) or under stabilising selection (B, C; mean(α=β)=45). A-B) False positive rate across 1% quantiles of ΔSVM. C) False positive rate as a function of the number of substitutions. D) False positive rate as a function of the fraction of the dataset analysed, from 100% of peaks down to the top 1%, ranked by derived SVM score (post-simulation binding affinity proxy). *N* = 10,000 simulated peaks; FDR ≤ 0.01.

Distinguishing stabilising selection from random drift is inherently difficult, as most substitutions have low predicted effects on TF binding affinity (Supplementary Figure 7). We therefore assessed RegEvol’s performance in detecting stabilising selection across a range of simulation parameters (Supplementary Figure 8). Moreover, because of the asymmetry of the DPE toward negative changes, the distribution of ΔSVM in random permutations becomes increasingly negative with the number of substitutions. Consequently, under stabilising selection, the false discovery rate of the permutation test positively correlates with sequence divergence and can reach a FPR of 0.05 with high number of substitutions (Figure 4.C, Spearman’s rho=0.96, p=7.4e-16). By integrating the ΔSVM distribution as a DPE for each peak and explicitly modeling stabilising selection around ΔSVM = 0, RegEvol effectively controls for this bias (FPR=0).

An ascertainment bias on ChIP-seq peaks has been suggested to affect permutation tests, as higher SVM score peaks are both more likely to be called and to show high positive ΔSVM relative to the ancestral sequence (Jiang and Zhang 2024). To test this, we subsampled peaks ranked by derived SVM score after random drift simulations, and assessed the proportion detected under directional selection (i.e FPR). RegEvol maintained a FPR close to 0 across all fractions of the dataset analysed (maximum FPR = 1e-4 for the 100% dataset). By contrast, the permutation test showed a clear increase in FPR as subsampling was enriched for higher SVM scores, reaching an FPR of 0.09 when only the top 1% of peaks was retained (Figure 4D). (Figure 4.D). Applying the same subsampling to the previously analysed human CEBPA dataset (Liu and Robinson-Rechavi 2020), the correlation for RegEvol remained low, with only the highest SVM category showing a slight increase, potentially reflecting a true biological signal (Supplementary Figure 9). These results demonstrate that RegEvol is robust to peak strength, supporting its reliability in real datasets where ChIP-seq signal intensity may influence peak calls.

To evaluate practical limitations of RegEvol, we examined how detection power is affected by the number of substitutions per regulatory element and multiple testing correction (Supplementary Figures 10-11). In empirical datasets, many peaks do not have enough substitutions, particularly in closely related species. For example, in mammalian datasets, the median number of substitutions per peak can be low depending on the divergence since the closest ancestral state (human –chimpanzee median substitution number = 2.8), which reduces statistical power and leads to a scarcity of low p-values after multiple testing correction (Supplementary Figure 11). In Drosophila, higher divergence results in more substitutions per peak, providing a more stable detection power across regulatory elements (Supplementary Figure 10). Given these observations and the low true positive rate for peaks with few substitutions even under strong simulated directional selection (Supplementary Figure 6), we restricted our analyses to peaks with at least four substitutions in vertebrate datasets (Supplementary Figure 12). This corresponds to an average of 1.7% sequence divergence (median length=291bp; Supplementary Figure 13-14).

### 5 -Analysis of *Drosophila melanogaster* ChIP-seq peaks

To illustrate the utility of RegEvol, we applied it to a large-scale dataset of transcription factor binding sites in *Drosophila melanogaster*. Using ChIP-seq peaks from the modERN consortium (Kudron et al. 2024), we analysed regulatory elements across multiple developmental stages and transcription factors (Supplementary Table 2). For each peak, we computed the DPE using gkm-SVM models trained on the corresponding TF and inferred the most likely selection regime since the divergence from *D. simulans*, based on ancestral sequences reconstructed from whole-genome alignments.

We inferred a total proportion of 5.1% of fruit fly peaks under directional selection (Figure 5A), out of 2.8M tested. This is not due to high permissiveness of the test, since on mammalian ChIP-seq peaks we find much lower proportions of directional selection (Supplementary Figure 12). This is consistent with reports of a higher proportion of amino acid substitutions fixed by directional selection, and generally higher efficiency of selection, in fruit flies than in mammals (Eyre-Walker 2006; Sella et al. 2009; Lin et al. 2025). Among the fly peaks, there is considerable variation between TFs and conditions, from almost none to 50% of peaks inferred under directional selection. With the low developmental resolution of those data, there is no strong trend of differences over ontogeny but there is less signal of directional selection in prepupal and pupal stages (Supplementary Figure 15).

**Figure 5.**
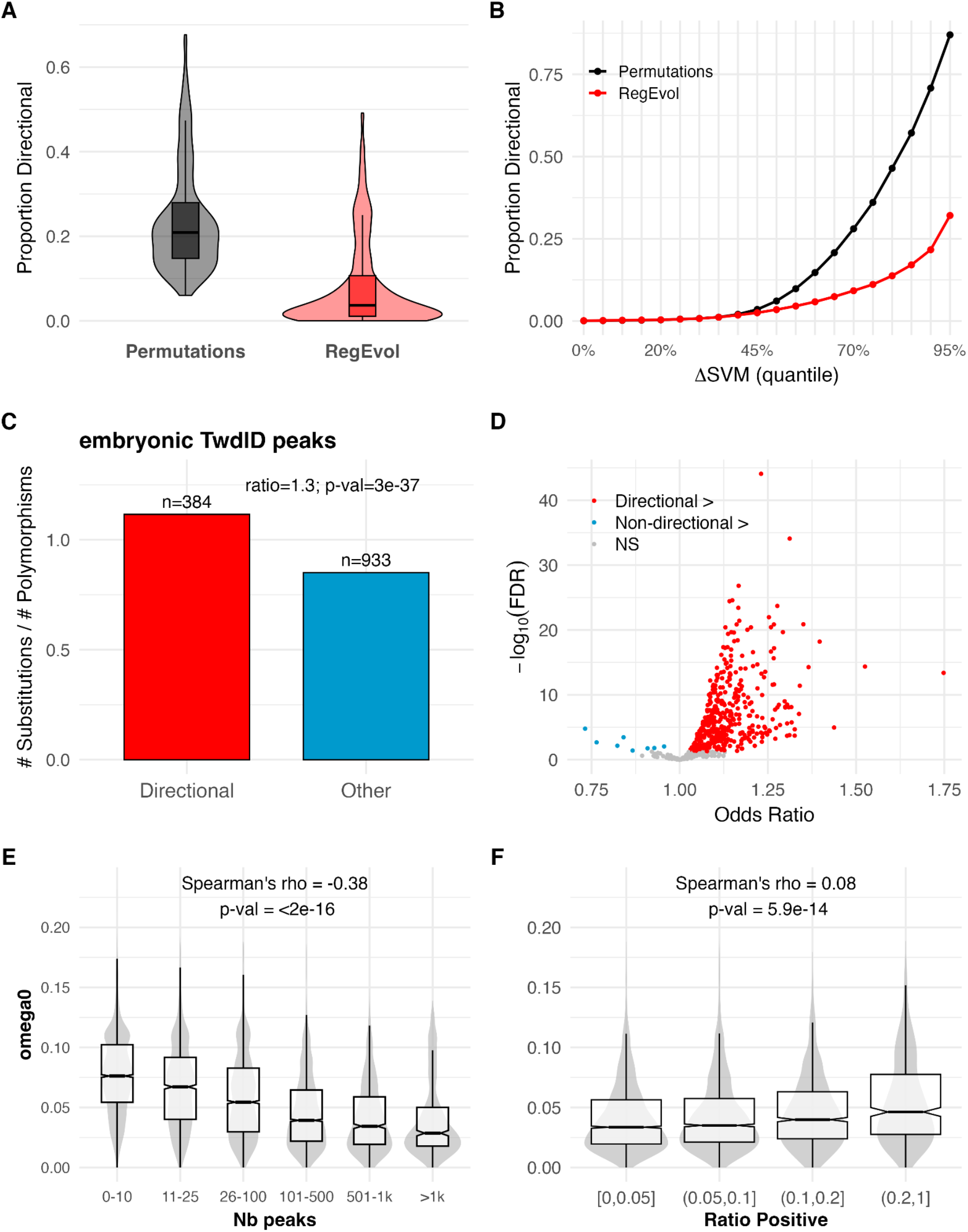
Application of RegEvol to Drosophila melanogaster transcription factor binding sites. **A.** Proportion of peaks inferred to be under directional selection using the original permutation test (black; *p* ≤ 0.01) and RegEvol (red; FDR ≤ 0.05). **B.** Proportion of peaks under directional selection across the 5% quantiles of derived ΔSVM values for all peaks. **C.** Ratio of substitutions to single-nucleotide polymorphisms (SNPs) in embryonic *TwdlD* peaks inferred by RegEvol under directional selection (red; *N* = 385) or not (blue; *N* = 877). The odds ratio and Fisher’s exact test between the two categories are indicated. **D.** Odds ratios between directional and non-directional 417 peaks across datasets (restricted to experiments with ≥10 directional peaks). Experiments where directional peaks show higher ratios (red; odds ratio > 1), lower ratios (blue; odds ratio ≤ 1), or no significant difference (grey; FDR > 0.05) are shown. **E-F.** Relationship between purifying selection on protein-coding genes (ω₀) and the total number of associated peaks (**E**) or the proportion of their peaks inferred under directional selection (**F**). Spearman’s correlation coefficients were computed across all genes.

As expected, the signal for directional selection increases with the ΔSVM (Figure 5B): peaks with larger predicted phenotypic effects are more often inferred to have evolved under directional selection. Among peaks under directional selection, the vast majority show selection to increase binding affinity (99.8%). This asymmetry is only partially explained by the shape of the DFE: the difference in detection power between negative and positive changes in simulations (Figure 3) is far smaller than the asymmetry observed empirically, suggesting a contribution from additional factors. Detection of directional selection towards reduced affinity is inherently limited by ChIP-seq sensitivity: a peak must retain sufficient binding affinity to be called, truncating the range of observable negative ΔSVM values. Accordingly, essentially the entire range of ΔSVM expected under positive directional selection falls within this detectable range, whereas only a small fraction of the range expected under negative directional selection does (overlap coefficient = 0.50 vs 0.16, Supplementary Figure 16). Moreover, the large effective population size of Drosophila is expected to make selection highly efficient, so that sequences evolving towards reduced affinity are quickly eliminated, leaving fewer sequences segregating long enough to be detected as evolving towards decreased affinity. We detect few peaks under stabilising selection, which can be explained by the framing of our test relative to the nature of purifying selection: stabilising selection tends to eliminate mutations, leading to few substitutions and a low power of the likelihood test. The role of stabilising selection in our model is essentially to avoid false positives in the test of directional selection; however, our approach is poorly adapted to inferring the proportion of peaks evolving under stabilising selection *per se*.

A key prediction of recent directional selection is a local reduction in polymorphism due to selective sweeps, together with increased inter-species divergence from fixation of mutations (McDonald and Kreitman 1991; Fu and Akey 2013). Accordingly, we expect a higher substitution-to-SNP ratio in peaks inferred to have evolved under directional selection. This expectation is confirmed in D. melanogaster, with almost all experiments supporting this pattern (74.8% experiment, odds ratio > 1 and Fisher’s exact test FDR ≤ 0.05) (Figure 5 C, D). Because our model gains power with increasing numbers of substitutions, this pattern could in principle reflect a bias. To evaluate this, we compared substitutions and SNPs separately and found that peaks under directional selection have both significantly more substitutions and fewer SNPs (Wilcoxon test p-values ≤ 5×10⁻⁴; Supplementary Figure 17), consistent with recent selective sweeps rather than a power artefact. This provides independent validation, as SNP frequencies are not used by the model, and shows that our k-mer-based DFE inference aligns with classical expectations of natural selection.

While it is difficult to reliably associate TF ChIP-seq peaks with the genes they regulate, the closest genes provide a good approximation, especially in compact genomes such as that of *D. melanogaster* (Kudron et al. 2024). As expected, protein-coding genes under stronger purifying selection (ω₀) are associated with more TF peaks (Figure 5E), thus presumably more complex regulation and more pleiotropy. There is also an association, although weaker, between selection on protein-coding genes and on regulatory sequences (Spearman’s rho=0.08, p-values=8e-16, Figure 5F): proteins under weaker purifying selection are associated with a higher proportion of peaks under directional selection. We do not find an association between adaptive selection on protein-coding genes (ω2) and the proportion of directional selection on their associated peaks.

Genes associated with peaks under directional selection are enriched for morphogenesis and developmental processes (Supplementary Figure 17). However, genes linked to a larger total number of peaks are also overrepresented in these functional categories. When considering the ratio of peaks under directional selection, no significant functional enrichment was observed (Supplementary Figure 17). The lack of functional enrichment may reflect a limitation of this approach. By aggregating TF peaks from 740 different experiments and contexts, the resulting gene set may be too large and functionally heterogeneous for conventional functional tests. To explore whether signals of directional selection could be detected at a broader scale, we applied TopAnat (Komljenovic et al. 2018a; Bastian et al. 2021), which identifies anatomical structures where a gene set is preferentially expressed. Genes with a high proportion of selected TF peaks (≥0.2) were enriched in reproductive tissues (testis, seminal fluid glands, ovary) and immune-associated organs (fat body, Malpighian tubule) (Supplementary Figure 18).

### 6 -Tissue-level aggregation reveals directional selection in human CTCF

Applying RegEvol to mammalian ChIP-seq peaks individually yielded a low proportion of elements classified under directional selection (Supplementary Figure 12). This pattern is inherent to lineage-specific evolutionary analyses conducted over short evolutionary distances, where the limited number of substitutions per regulatory element constrains statistical power. To address this limitation, we performed a tissue-level aggregation analysis inspired by the SUMSTAT framework of Daub et al. (2017). For each tissue, we computed cumulative SUM scores from the differences between the likelihood of the Directional model and the Neutral model across active peaks (Material and Methods). Analyses were controlled for sample size through resampling to ensure comparability across tissues. We applied this strategy to human CTCF ChIP-seq peaks detected across multiple tissues (ENCODE Project Consortium 2012).

SUM scores displayed a broad plateau across most tissues, followed by a clear upward shift culminating in a distinct cluster of high-ranking tissues (Figure 6). The highest-ranked tissues were predominantly central nervous system–associated cell types together with male reproductive tissues, whereas peripheral nervous tissue and the majority of other somatic tissues remained within the lower plateau. Formally grouping tissues by biological system revealed that both the Nervous system and the Male reproductive system exhibited higher cumulative SUM scores compared to all other systems (Wilcoxon rank-sum tests between systems significant, p < 2.2e-16; Figure 6). Effect size estimates confirmed a strong positive shift, i.e. excess of positive selection signal, for the Nervous system (Cliff’s δ = 0.55) and a smaller positive shift for the Male reproductive system (Cliff’s δ = 0.23). No other systems had positive shifts (all Cliff δ and Wilcoxon p-values in Supplementary Table 3), including immune-associated tissues. A similar analysis performed on chimpanzee homologous sequences of these CTCF peaks recovered a comparable but attenuated enrichment for the Nervous system (Cliff’s δ = 0.47) and a weaker signal for the Male reproductive system (Cliff’s δ = 0.12; Supplementary Figure 21). This is consistent with a common acceleration of regulatory sequence evolution across primates (Zhuang et al. 2023). The weaker signal in chimpanzee could be due to the use of human experimental peaks, or to increased selection along the human lineage.

**Figure 6.**
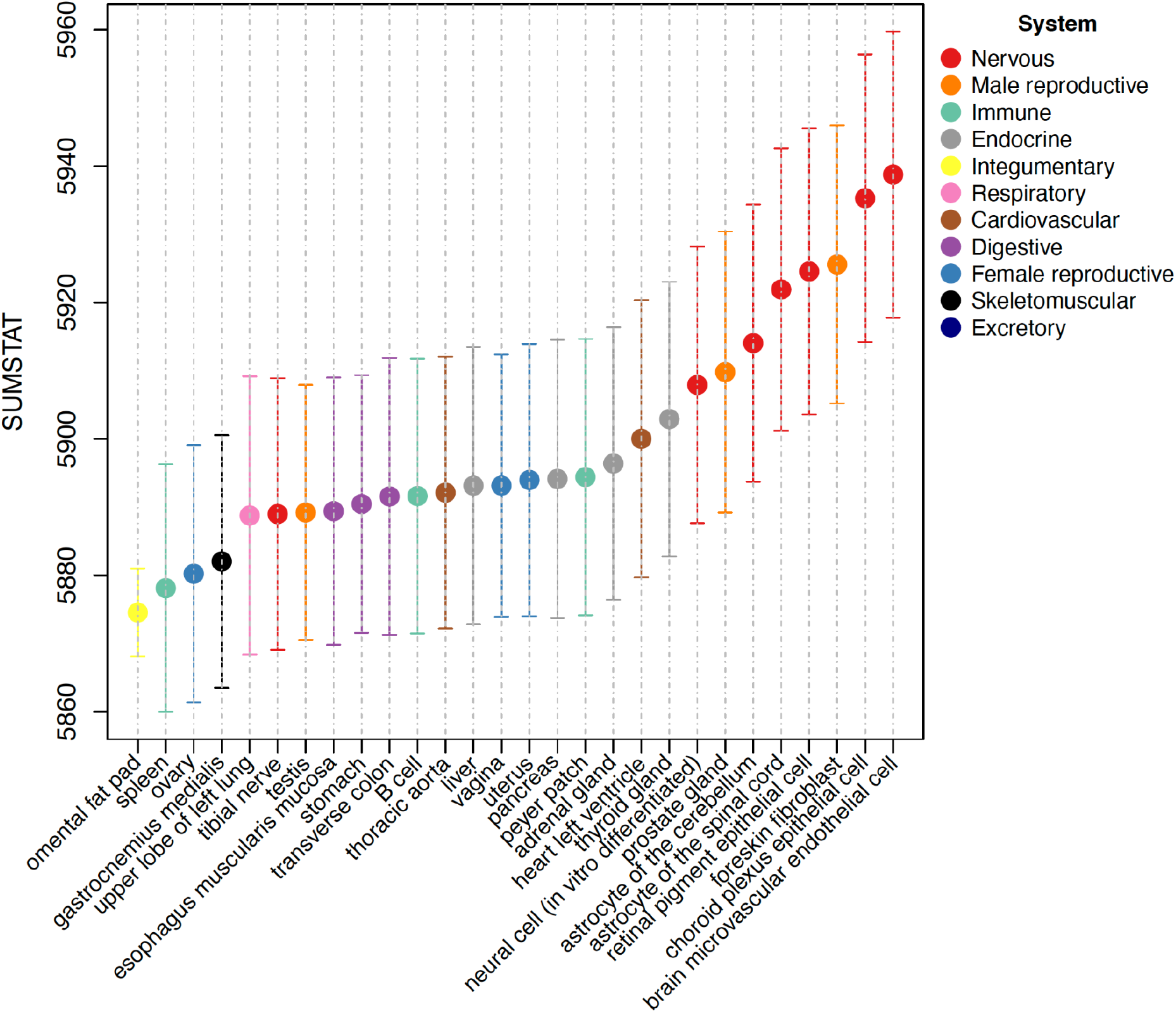
Tissue-level aggregation of directional selection on human CTCF peaks. Median SUMSTAT scores across 10,000 resamplings of 5,000 human CTCF peaks per tissue. Bars indicate the standard deviation across resamplings. SUMSTAT scores were calculated following the framework from Daub et al. (2017) by taking the fourth-root of the per-peak Δlog-likelihood (Directional minus Neutral model) and summing across all peaks within each tissue. Tissues are sorted by median SUMSTAT and colored according to organ system.

## Discussion

Understanding how regulatory sequences evolve is essential for linking genotype to phenotype, yet most evolutionary analyses of non-coding DNA still rely on substitution rates or sequence conservation (Adam Siepel et al. 2005; Pollard et al. 2010). These approaches are powerful for identifying constraints but often lack sensitivity to detect selection acting on specific molecular functions (Ward and Kellis 2012; Villar et al. 2015), especially directional selection. RegEvol addresses this limitation by testing how predicted mutational effects on transcription factor (TF) binding have been shaped by selection. By linking quantitative predictions of regulatory activity to patterns of sequence divergence, RegEvol provides a functionally informed and mechanistically interpretable view of regulatory evolution. In contrast to motif-centric models, the predictive component of RegEvol is trained on entire ChIP-seq peaks, capturing local sequence context and potential co-factors that influence TF binding (J. Yan et al. 2021). This shift from rate-based to molecular phenotype-based inference enables the detection of subtle selection signals that may not produce strong shifts in sequence conservation but shape regulatory activity. Moreover, RegEvol leverages causal role functional information to infer selective effect.

RegEvol builds on a previous permutation-based framework (Liu and Robinson-Rechavi 2020), which evaluated selection by comparing the summed ΔSVM of observed substitutions to a null distribution derived from random permutations. While this approach provided a useful baseline, it was sensitive to extreme substitution effects, lacked robust multiple-test correction, and was susceptible to ascertainment bias (Jiang and Zhang 2024). RegEvol addresses these limitations by modelling three nested selective regimes (neutral, stabilising, and directional), through fitness functions that link predicted TF-binding effects to substitutions patterns. Using a maximum-likelihood framework, our method identifies the selective regime that best explains the observed substitutions. Moreover, because inference is analytical rather than simulation-based, p-values remain well calibrated and support rigorous multiple-test correction, overcoming a key weakness of the permutation approach (Jiang and Zhang 2024). A key strength of RegEvol is its nested modeling of selective regimes, allowing direct likelihood-based comparison of directional and stabilising selection. This is important because the expectation under stabilising selection is centred on no phenotypic change (ΔSVM ≈ 0), whereas the permutation-based null is typically slightly negatively shifted. This asymmetry has little impact when few substitutions are available but leads to a marked inflation of false positives with increasing divergence (Figure 4). Including a stabilising regime corrects this bias and improves robustness by providing an intermediate model between neutrality and directional selection.

RegEvol remains robust across a wide range of divergence levels. Simulations show that it is conservative at short timescales, minimizing false positives, and gains power while maintaining specificity as substitutions accumulate. Even when the overall signal is modest, RegEvol detects directional selection when substitution effects are consistent. By modelling the full distribution of predicted effects rather than focusing on extreme values, the method emphasizes consistency over magnitude, reducing false positives driven by a few large-effect mutations and nearly eliminating ascertainment bias for the detection of directional selection towards increased binding affinity. This yields conservative inference, a desirable property when testing for positive selection (Jordan and Goldman 2012; Venkat et al. 2018; Wisotsky et al. 2020; Soni et al. 2023). A small number of strong-effect substitutions may therefore be insufficient to support directional selection, reflecting a trade-off between robustness and sensitivity. In contrast, a permutation-based approach can interpret extreme mutations as evidence of selection, potentially increasing power in rare-event scenarios but at the cost of elevated false-positive rates.

By decoupling substitution rate from phenotypic effect, RegEvol provides a functionally informed view of regulatory evolution, analogous to how coding sequence evolution can be interpreted through the nature of amino acid changes rather than their frequency (Halpern and Bruno 1998; Rodrigue et al. 2021). While coding sequences are constrained by protein structure and pleiotropy, regulatory elements evolve under different pressures, including redundancy, turnover, and modularity, which enhance evolvability but complicate evolutionary analysis (Wittkopp and Kalay 2012; Kim and Wysocka 2023). Application to *Drosophila melanogaster* ChIP-seq data illustrates this framework empirically: 5.1% of peaks were inferred under directional selection, consistent with previous reports of adaptive evolution in fly transcription factor binding sites (He et al. 2011) and with the higher efficiency of selection in flies compared to mammals (Eyre-Walker 2006; Sella et al. 2009; Lin et al. 2025). Peaks under directional selection showed elevated substitution-to-SNP ratios, consistent with recent selective sweeps (McDonald and Kreitman 1991; Fu and Akey 2013). These peaks were also associated with genes enriched in reproductive and immune tissues, systems that are major targets of sexual selection and host–pathogen interactions and are known to evolve rapidly (Buchon et al. 2014; Vaibhvi et al. 2022), supporting the biological relevance of the inference. This framework thus provides a link between regulatory sequence evolution and downstream phenotypic effects, offering new perspectives on the co-evolution of regulatory elements and associated genes, and more broadly connecting regulatory variation to organismal adaptation.

Reduced power under low substitution numbers is an inherent limitation of lineage-specific tests. In some regions, this scarcity of substitutions may result from strong purifying selection, in which case conservation-based methods complement rather than compete with functionally informed approaches such as RegEvol. In the mammalian dataset, the limited number of substitutions per regulatory element constrained per-peak inference, motivating the use of a tissue-level aggregation strategy. We applied this approach to human CTCF binding sites, summing per-peak likelihood differences between directional and neutral models across biologically coherent tissue groups, following principles similar to SUMSTAT analyses of coding sequences (Daub et al. 2017). Aggregating likelihoods preserves the contribution of individual peaks and avoids composite sequences, revealing coherent enrichment in nervous and male reproductive tissues, consistent with previous evidence of accelerated and adaptive evolution in these systems (Dorus et al. 2004; Khaitovich et al. 2006; Liu and Robinson-Rechavi 2020). While we focused on tissue-level aggregation here, the same framework could be extended to other biologically meaningful sets, such as pathways, cell types, or regulatory modules, allowing RegEvol to uncover directional trends that are otherwise undetectable at the level of individual elements.

Detection of directional selection towards decreased affinity, however, remains fundamentally limited by the truncated nature of the data itself: because ChIP-seq peaks require sufficiently high binding affinity to be called in the first place, sequences that have already evolved strongly reduced affinity are inherently absent from the dataset, a limitation that no statistical method can overcome. As a result, our estimates of the prevalence of directional selection towards decreased affinity are conservative and likely underestimate its true extent.

Improving the genotype-to-phenotype map remains essential to extend RegEvol’s reach. While gkm-SVM models remain interpretable and TF-specific, capturing the additive effects of individual substitutions, they have limited ability to model complex regulatory features such as motif syntax, long-range dependencies, or non-additive interactions (de Boer and Taipale 2024). Recent deep learning models trained on high-resolution functional genomics data have shown the ability to learn these features directly from sequence, providing more accurate, context-aware predictions of mutational impact (Avsec et al. 2021; Benegas et al. 2023; Karollus et al. 2024). Because RegEvol is agnostic to the underlying predictive model, it can incorporate these and future advances, producing more informative phenotypic effect distributions and improving sensitivity in complex regulatory landscapes. Moreover, regulatory elements often integrate inputs from multiple TFs, chromatin states, and cellular contexts, and the functional impact of a mutation may vary across these layers (Hu and Tee 2017; Kurafeiski et al. 2018). Probabilistic frameworks, such as Bayesian or mixed-effect models, could capture heterogeneous and overlapping selective pressures within a single element, capturing this complexity more realistically than a single fitness landscape per element.

Together, these developments position RegEvol as a flexible and extensible framework for studying the evolution of regulatory sequences in a functionally informed manner. By integrating predictive models of molecular phenotypes, which link genotype to quantitative regulatory effects, with well-established evolutionary equations that map phenotypes to fitness, RegEvol provides a principled approach to detect directional selection, complementing traditional conservation-based analyses. As functional genomics, predictive modeling, and comparative data continue to advance, including the ability to transfer predictive models across species, RegEvol could support more systematic, multi-lineage analyses. This would help bridge the gap between regulatory sequence variation and phenotypic evolution and offer new insights into the molecular mechanisms underlying adaptation across evolutionary timescales.

## Materials and Methods

### Catalogue of Transcription Factor Binding Sites

We used pre-processed ChIP-seq peak data for *Drosophila melanogaster* TFBS from the modENCODE (model organism Encyclopedia of DNA Elements) and modERN (model organism Encyclopedia of Regulatory Networks) consortia (Kudron et al. 2024). It provides a harmonised dataset across 740 experiments covering 645 TFs and five developmental stages (Supplementary Table 2).

### ChiP-seq peaks detection and coverage quantification

We collected publicly available ChIP-seq data for five transcription factors (TF) across ten species: *Homo sapiens*, *Macaca macaca*, *Mus musculus*, *Mus spretus*, *Mus caroli*, *Rattus norvegicus*, *Canis lupus familiaris*, *Felis catus*, *Oryctolagus cuniculus, Gallus gallus*, *Drosophila melanogaster* (Ballester et al. 2014; D. Schmidt et al. 2010; Dominic Schmidt et al. 2012; Rensch et al. 2016; Stefflova et al. 2013) (Supplementary Table 1). To ensure consistency across datasets, we re-processed all raw sequencing data using the NextFlow ChIP-seq pipeline v2.0 (Ewels et al. 2022), specifying Bowtie2 (Langmead and Salzberg 2012) as the aligner and using the narrow_peak option for peak calling with MACS2 (Y. Zhang et al. 2008). For downstream analyses, we retained only peaks located on complete chromosomes, excluding unplaced scaffolds.

Read coverage was computed at each base within ChIP-seq peaks using bamCoverage, normalised by counts per million (Ramírez et al. 2014). For each experiment, we calculated the consensus peak coverage by summing normalised read counts across replicates. Peak summits were defined as the position with the highest normalised read coverage within each peak. In cases where multiple positions shared the maximum value, we selected the position with the highest mean coverage across a 10 bp window centred on the candidate site.

### Whole genome alignment and ancestral state

We retrieved publicly available whole-genome alignments generated with Progressive Cactus. For *Drosophila melanogaster*, we used a 20-species alignment (Peng and Zhao 2024), and for vertebrates, we used the 241-species alignment from the Zoonomia Consortium (Armstrong et al. 2020). These alignments include both reference genomes and inferred ancestral genomes at each internal node of the species tree.

For each analysed species, we used the reference genome as the query and extracted alignments with the closest related species and their most recent common ancestor. Aligned sequences were retrieved using the hal2maf tool from the Comparative Genomics Toolkit (Hickey et al. 2013), with the noDupes option enabled to exclude putative duplicated regions. We extracted the alignment of each set of ChIP-seq peaks using their genomic coordinates as input for the mafsInRegion tool from UCSC (Nassar et al. 2023).

### gkm-SVM model training and validation

To model the relationship between DNA sequence and TF binding affinity, we trained gapped k-mer support vector machine (gkm-SVM; Ghandi et al. 2014) models for each TF and tissue. The positive training set consisted of genomic regions identified as ChIP-seq peaks in the reference genome. The negative set was generated by randomly sampling genomic sequences matched for length, GC content, and repeat content using the genNullSeqs function from the gkmSVM R package (Ghandi et al. 2016). When pre-built BSgenome objects were not available for the genome assemblies analysed, we constructed them following the BSgenome R package guidelines (Pagès 2024), using annotations from UCSC and GenBank (Supplementary Table 1).

Each model was trained using the gkmtrain function from the LS-GKM software (D. Lee 2016), with default parameters except for a k-mer length of 10 bp. Model performance was evaluated using 5-fold cross-validation using the –x 5 option. After training, we used the gkmpredict function to compute SVM weights for all possible 10-mers. Experiments with fewer than 1,000 peaks or with an associated model AUC below 0.8 were excluded from downstream analyses (Supplementary Figure 19).

### SVM Score Computation for Peaks and Positions

We computed SVM scores for each ChIP-seq peak at two levels. At the peak level, the total SVM score was calculated as the sum of SVM weights from a 10 bp sliding window with a 1 bp step size. At the positional level, the SVM score for a specific nucleotide position was defined as the sum of SVM weights for all overlapping windows that include that position. These two levels of scoring were used for downstream analyses of both peak-level and site-specific effects.

### In Silico Mutagenesis and ΔSVM Computation

For each ChIP-seq peak, we performed in silico mutagenesis by introducing all possible single-nucleotide mutations. The change in predicted binding affinity for each mutation was computed as ΔSVM, defined as the difference in SVM score between the mutated and ancestral sequence. This yielded a Distribution of Phenotypic Effects (DPE) for each peak, representing the genotype-to-phenotype landscape.

Observed substitutions were identified by comparing the reference sequence to its inferred ancestral state. Indels were removed using TrimAl with the “nogaps” option (Capella-Gutiérrez et al. 2009). SVM scores were computed for both the reference and ancestral sequences using the same sliding window approach, and ΔSVM values were calculated for each observed substitution.

### Substitution matrix inference

To account for variation in mutation and substitution rates across the genome, we estimated substitution matrices for each chromosome. Whole-genome alignments were split by chromosome using the mafSplit tool, and exonic regions were masked using GFF annotations. We then ran PhyML (Guindon et al. 2010) on each chromosome alignment using a Generalised Time-Reversible model with the params=r option to optimise substitution rate parameters. These matrices were used to weight mutation probabilities in the evolutionary model.

### Evolutionary Model Inference via Likelihood Optimisation

To infer the selection regime acting on each ChIP-seq peak, we modeled the evolutionary process using three nested phenotype-to-fitness maps parameterized by Beta distributions: (i) a neutral model with a flat fitness landscape (α = β = 1), (ii) a stabilising selection model with maximal fitness at the ancestral phenotype (α = β ≠ 1), and (iii) a directional selection model allowing asymmetric fitness landscapes (α ≠ β).

For each possible single-nucleotide mutation within a peak, we computed the predicted phenotypic effect (ΔSVM), the corresponding selection coefficient (as the log-ratio of fitness between derived and ancestral phenotypes), and the fixation probability under an origin-fixation model in which a mutation’s probability of fixation depends on its selection coefficient. These were combined with mutation probabilities derived from chromosome-specific substitution matrices to calculate a substitution probability for every in silico mutation.

The likelihood of the observed substitutions was then computed by summing the log-probabilities of the corresponding mutations under each model. Model parameters (α, β) were optimised using the scipy.optimize.minimize function (Virtanen et al. 2020). To compare the performance of each model and determine the most likely evolutionary scenario for each peak, we performed likelihood ratio tests using a p-value threshold of 1%. A detailed mathematical formalism of this framework is provided in Supplementary Materials.

### Improvement on the Permutation Test

We refined the previously published Permutation Test to improve accuracy and interpretability (Liu and Robinson-Rechavi 2020). Mutation and substitution rates are now incorporated using chromosome-specific substitution matrices. Substitution positions are sampled without replacement to match the observed count, avoiding undersampling artefacts. We also implemented a two-sided test to detect both increases and decreases in binding affinity, and applied false discovery rate (FDR) correction to account for multiple testing.

### Transcription factor binding motif recovery

To assess whether the gkm-SVM models captured known TF binding preferences, we extracted the SVM weights for all possible 10-mers from the trained models for human and mouse datasets. We selected the top 100 scoring 10-mers for each model and aligned them using Clustal Omega with default parameters (Sievers et al. 2011). The resulting alignments were submitted to the TomTom motif comparison tool (Gupta et al. 2007) in the MEME Suite website (Timothy L. Bailey et al. 2015) to identify matches against known TF motifs in the JASPAR CORE 2022 database (Castro-Mondragon et al. 2022).

### Sequence conservation metrics

To compare model-based predictions with evolutionary conservation, we retrieved per-base conservation scores from the UCSC Genome Browser (Perez et al. 2025). Specifically, we used phastCons (Siepel et al. 2005) and phyloP scores (Pollard et al. 2010) from the 17-way vertebrate alignment for human, the 60-way glire subset for mouse, and the 27-way alignment for *Drosophila melanogaster*. For each ChIP-seq peak, we computed the conservation score at every nucleotide position and summarised the signal by calculating the average score across the peak.

### Simulation of sequence evolution

To assess the accuracy and robustness of our selection inference framework, we simulated the evolution of regulatory sequences under controlled evolutionary scenarios. From randomly sampled reconstructed ancestral sequences of drosophila CTCF ChIP-seq peak, we computed the predicted phenotypic effect (ΔSVM values) for all possible single-nucleotide mutations, along with mutation probabilities derived from chromosome-specific substitution matrices.

Substitutions in these sequences were sampled probabilistically according to their mutation rate, predicted phenotypic effect, and fixation probability under a specified evolutionary model. These models included neutral evolution, stabilising selection around the ancestral phenotype, and directional selection favouring increased or decreased binding affinity. Each model was parameterised using Beta distributions to define the shape and strength of the phenotype-to-fitness map.

Simulations were performed across a range of conditions, varying the number of substitutions per sequence, the strength of selection and the proportion of neutral mutations. This allowed us to generate synthetic datasets reflecting different evolutionary regimes, which were then used to benchmark the performance of RegEvol and the Permutations Test.

### Drosophila divergence and polymorphism

Polymorphism data from the *Drosophila melanogaster* Genetic Reference Panel (DGRP) (Mackay et al. 2012) were retrieved and lifted over from the dm3 to the dm6 genome assembly using LiftOver (Hinrichs 2006). Overlap with TF peaks was determined using bedtools intersect. Fixed substitutions between *D. melanogaster* and *D. simulans* were counted within each peak, and a substitution-to-SNP ratio was calculated for every peak. Fisher’s exact tests were then applied for each experiment to compare peaks inferred to have evolved under directional selection against all other peaks.

### Gene Ontology and Anatomical Enrichments

Associations between ChIP-seq peaks and genes were obtained from the modERN consortium based on genomic proximity (Kudron et al. 2024). Gene Ontology enrichment analysis was performed using the *clusterProfiler* R package (version 4.16) with gene annotations from *org.Dm.eg.db* (version 3.21.0) for *Drosophila melanogaster* (Wu et al. 2021). Enrichment was computed against a background of all genes with at least one associated peak. We tested genes with at least one peak under directional selection, ranked gene lists based on either the total number of peaks or the number of peaks under directional selection, and genes with a directional-to-total peak ratio above 0.25 compared to all genes with at least two associated peaks.

Anatomical enrichment was carried out using TopAnat in the BgeeDB R package (version 2.34.0) with Bgee release 15.2 (Komljenovic et al. 2018; Bastian et al. 2025). TopAnat identifies anatomical entities where a gene set is preferentially expressed based on expression calls integrated by Bgee across anatomical structures. The analysis was restricted to RNA-seq expression calls and limited to *Drosophila melanogaster* fully formed stage (UBERON:0000066). Only the gene set with a directional-to-total peak ratio above 0.25 and the corresponding background with at least two peaks was tested. Enrichment was assessed using the *weight01* algorithm (Alexa et al. 2006), and anatomical entities with FDR ≤ 0.01 were considered significant.

### Drosophila coding sequences evolution

To assess whether genes associated with TF peaks also show signatures of coding sequence evolution, we retrieved ω₀ and ω₂ estimates from Selectome (Moretti et al. 2014). Selectome applies branch-site codon models (Davydov et al. 2019) to orthologous gene alignments to estimate the strength of purifying selection (ω₀) and positive selection (ω₂) on specific phylogenetic branches. These values were extracted for *D. melanogaster* genes associated with TF peaks and used to test for correlations with the number of peaks and the directional-to-total peak ratio.

### Tissue-level aggregation analysis of human CTCF binding sites

We analysed ChIP-seq peak data for CTCF in *Homo sapiens* identified by the ENCODE consortium and pre-processed in a previous study (ENCODE Project Consortium 2012; Liu and Robinson-Rechavi 2020). This dataset contains merged CTCF peaks across multiple tissues and cell types, allowing peaks to be assigned to broad physiological systems. RegEvol was applied to each peak to estimate the likelihood of each selective model.

To increase statistical power and test for coordinated signals of directional selection across biologically related elements, we implemented a tissue-level aggregation analysis inspired by the SUMSTAT framework of Daub et al. (2017). For each peak *i*, we computed the log-likelihood difference (Λ_log_ *L_i_*) between two scenarios: first the maximum-likelihood (*L*) of the observed data *O_i_* derived under a scenario of directional selection, with α-β-are the two parameters of directional selection estimated at the maximum (see Supplementary Materials for detailed equations on the computation for the likelihood), and second the likelihood for a scenario of no selection *L*[*O_i_*], as:

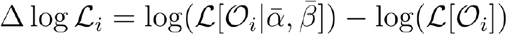

For each tissue, we performed 10,000 resamplings of 5,000 peaks to control for differences in sample size across tissues. For each resample, a cumulative SUM score was calculated as: Median and standard deviation of these SUM scores were then recorded for each tissue. Statistical significance of tissue-specific enrichment was assessed by comparing the distribution of SUM scores from each tissue to all other tissues using a Wilcoxon rank-sum test. Effect sizes were estimated using Cliff’s delta. This approach allows weak but concordant signals of directional selection to accumulate and be detected at the level of biologically coherent groups.

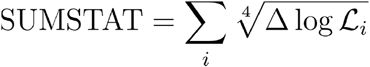

### Computation and Statistics

All data analyses were performed using R (version 4.5.0), Python (version 3.10), and Snakemake (version 7.24.0) for workflow management. The analyses were conducted on the high-performance computing infrastructure of the DCSR at the University of Lausanne. Multiple testing correction was applied using the Benjamini-Hochberg procedure to control the false discovery rate (FDR) across genomic regions. To assess potential ascertainment bias, we stratified ChIP-seq peaks by their SVM scores and evaluated detection rates across quantiles.

### Data Availability

The RegEvol pipeline, along with all scripts used to generate the analyses and figures, is available at the following stable GitHub URL: https://github.com/mrrlab/RegEvol. Additionally, all RegEvol results and data generated for this study are publicly available at the Zenodo repository: https://zenodo.org/records/18379807.

### Author Contributions

M.R-R and A.L. originally conceived the project, A.L. and T.L. designed and implemented the method and computational framework. A.L. conducted all analyses and wrote the initial draft of the manuscript. All authors discussed the results and contributed to the final version. M.R-R supervised the project.

### Funding

This work was supported by the Swiss National Science Foundation grant 207853 to Marc Robinson-Rechavi and grant 219757 to Nicolas Salamin.

