## Supplementary figures for "RegEvol: detection of directional selection in regulatory sequences through phenotypic predictions and phenotype-to-fitness functions"

**Supplementary Figure 1: gkm-SVM prediction along ChIP-seq peaks. 2**
**Supplementary Figure 2: Effect of peak quality on base-level SVM–ChIP-seq correlations. 3**
**Supplementary Figure 3: Correlation of gkm-SVM models across transcription factors and species. 4**
**Supplementary Figure 4: Principal component analysis of gkm-SVM models. 5**
**Supplementary Figure 5: Performance of RegEvol on simulated Drosophila CTCF ChIP-seq peaks. 6**
**Supplementary Figure 6: Detection performance of RegEvol under strong directional selection. 7**
**Supplementary Figure 7: Distribution of ΔSVM from all possible point mutations in CTCF peaks 8**
**Supplementary Figure 8: Detection performance of RegEvol under stabilising selection. 9**
**Supplementary Figure 9: Robustness to ascertainment bias in human data. 10**
**Supplementary Figure 10: Statistical behaviour of RegEvol on Drosophila melanogaster CTCF ChIP-seq peaks. 11**
**Supplementary Figure 11: Statistical behaviour of RegEvol on Homo sapiens CEBPA ChIP-seq peaks. 12**
**Supplementary Figure 12: Detection of directional selection by RegEvol across vertebrate species for five transcription factors. 13**
**Supplementary Figure 13: Characteristics of ChIP-seq peaks across species for five transcription factors 14**
**Supplementary Figure 14: Characteristics of Drosophila melanogaster modERN peaks. 15**
**Supplementary Figure 15: Drosophila melanogaster modERN peaks per developmental stage. 16**
**Supplementary Figure 16: Detection limits of negative directional selection. 17**
**Supplementary Figure 17: Drosophila melanogaster substitutions vs SNPs. 18**
**Supplementary Figure 18: Gene Ontology Enrichment. 19**
**Supplementary Figure 19: TopAnat Drosophila 20**
**Supplementary Figure 20: Model performance of Drosophila modERN experiments. 21**
**Supplementary Figure 21: Tissue-level aggregation of directional selection on chimpanzee homologs of human CTCF peaks. 22**

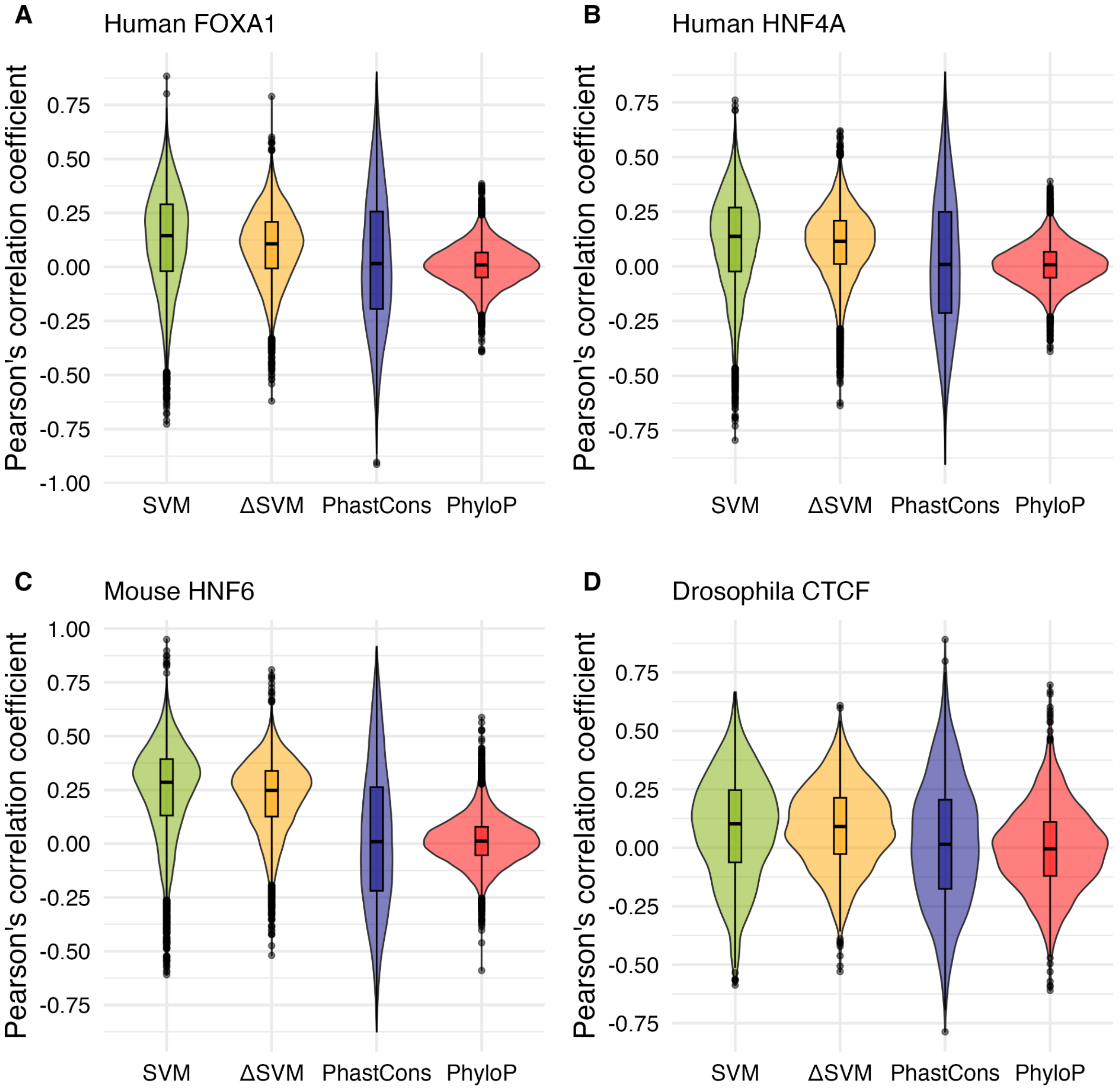


**Supplementary Figure 1: gkm-SVM prediction along ChIP-seq peaks.**
Distribution of Pearson correlation coefficients computed per peak between base-resolution ChIP-seq read coverage and four sequence-based metrics: SVM score, ΔSVM (change in SVM score from the ancestral state), phastCons, and phyloP. Correlations were calculated across nucleotide positions within each peak. Only ungapped positions were retained to ensure comparability across metrics. Panels show results for A) human FOXA1 (N = 14,420 peaks), B) human HNF4A (N = 18,503), C) mouse HNF6 (N = 17,493), and D) *Drosophila* CTCF (N = 2,309). All coverage normalisation, model training, conservation tracks, and filtering procedures are described in the Materials and Methods.

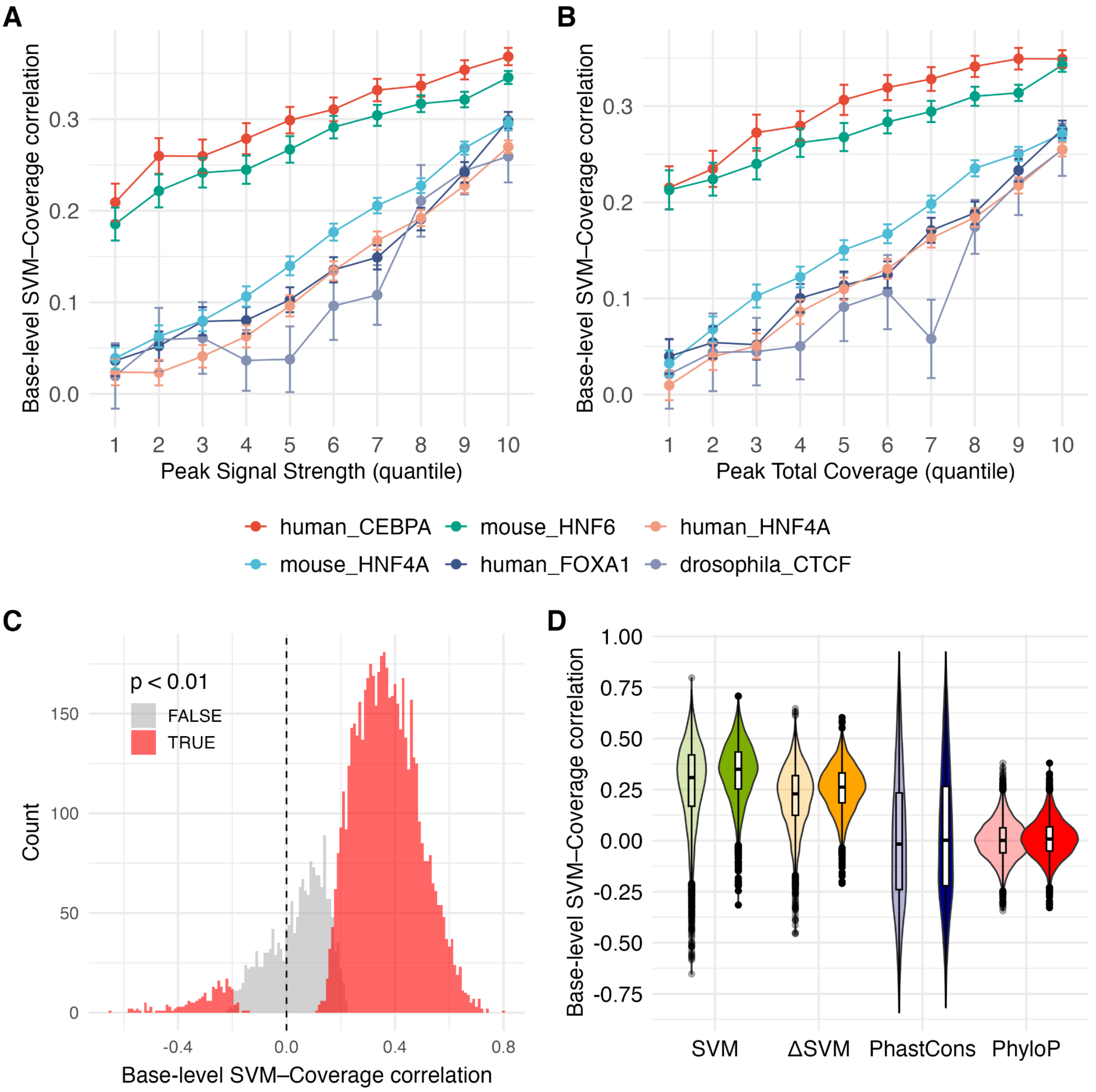


**Supplementary Figure 2: Effect of peak quality on base-level SVM–ChIP-seq correlations.**

For each peak, the Pearson correlation was computed between the SVM score and base-level ChIP-seq read coverage. Peaks were stratified by two quality metrics: **A)** signal strength (fold enrichment of ChIP over input) and **B)** total coverage, defined as the sum of reads across the peak. Peaks were binned into quantile for each metric. Points show median correlation, with error bars indicating approximate 95% confidence interval. Colours indicate different datasets. Across datasets, higher-quality peaks show significantly stronger base-level correlations (Pearson $r \approx 0.23–0.39$ for signal strength; 0.18–0.34 for total coverage). **C)** Distribution of per-peak Pearson correlation coefficients for the human CEBPA dataset, separated by statistical significance ($p < 0.01$ in red; not significant in grey). Most low or negative correlations are not statistically significant. **D)** Distribution of Pearson correlation coefficients between each metric (SVM, ΔSVM, phastCons, phyloP) and base-level read coverage across human CEBPA peaks. Two groups are shown: all peaks (light colors) and higher-quality peaks defined as the top 50% for both signal strength and total coverage (dark colors).

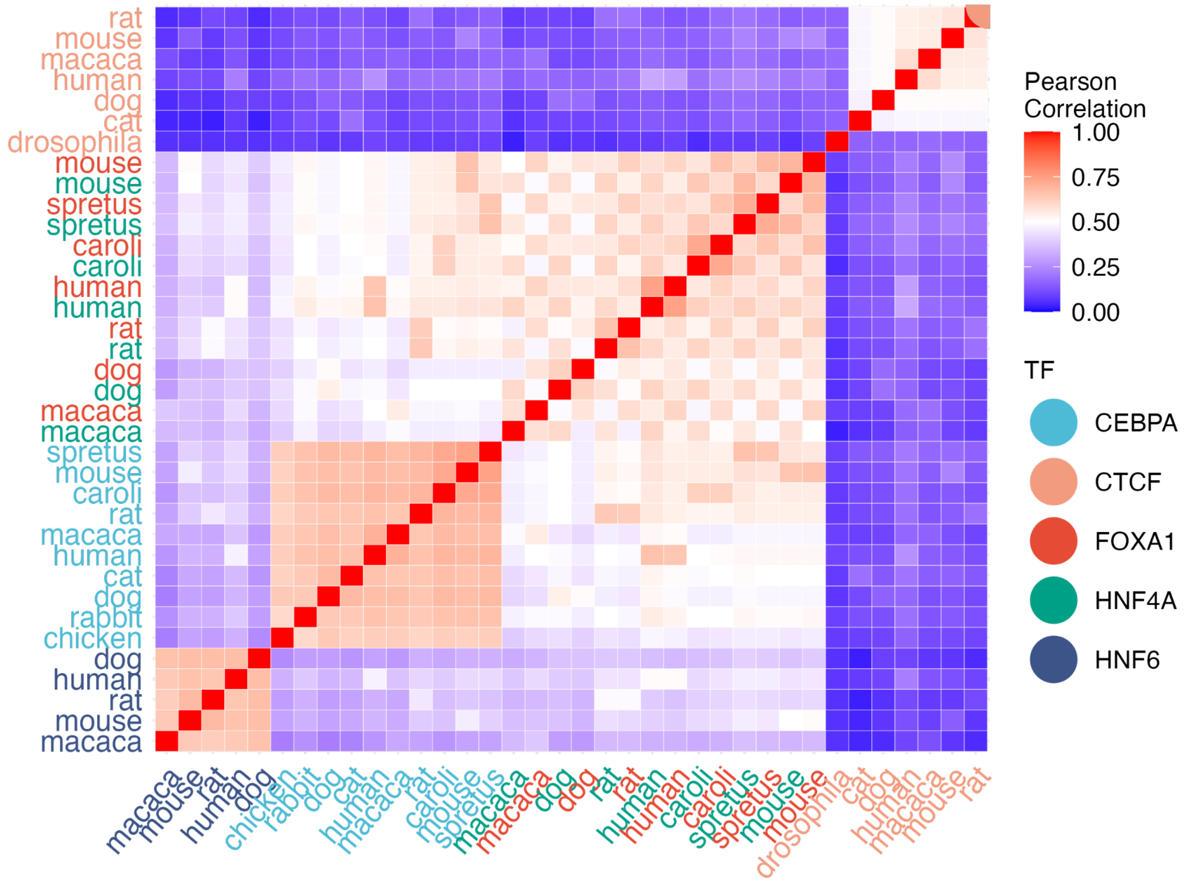


**Supplementary Figure 3: Correlation of gkm-SVM models across transcription factors and species.**

Pairwise Pearson correlations between gkm-SVM models trained on ChIP-seq peaks for five transcription factors across ten species (eight mammals, chicken, and *Drosophila*). Correlations were computed using SVM scores assigned to all possible 10-mers. Datasets are clustered by hierarchical clustering using correlation distance, reflecting similarity of the trained models. Axes are labelled by species, and text colour indicates the corresponding transcription factor: CEBPA (light blue), CTCF (orange), FOXA1 (red), HNF4A (green), HNF6 (blue). All correlations are positive, with models clustering primarily by transcription factor rather than species, consistent with TF-specific sequence preferences.

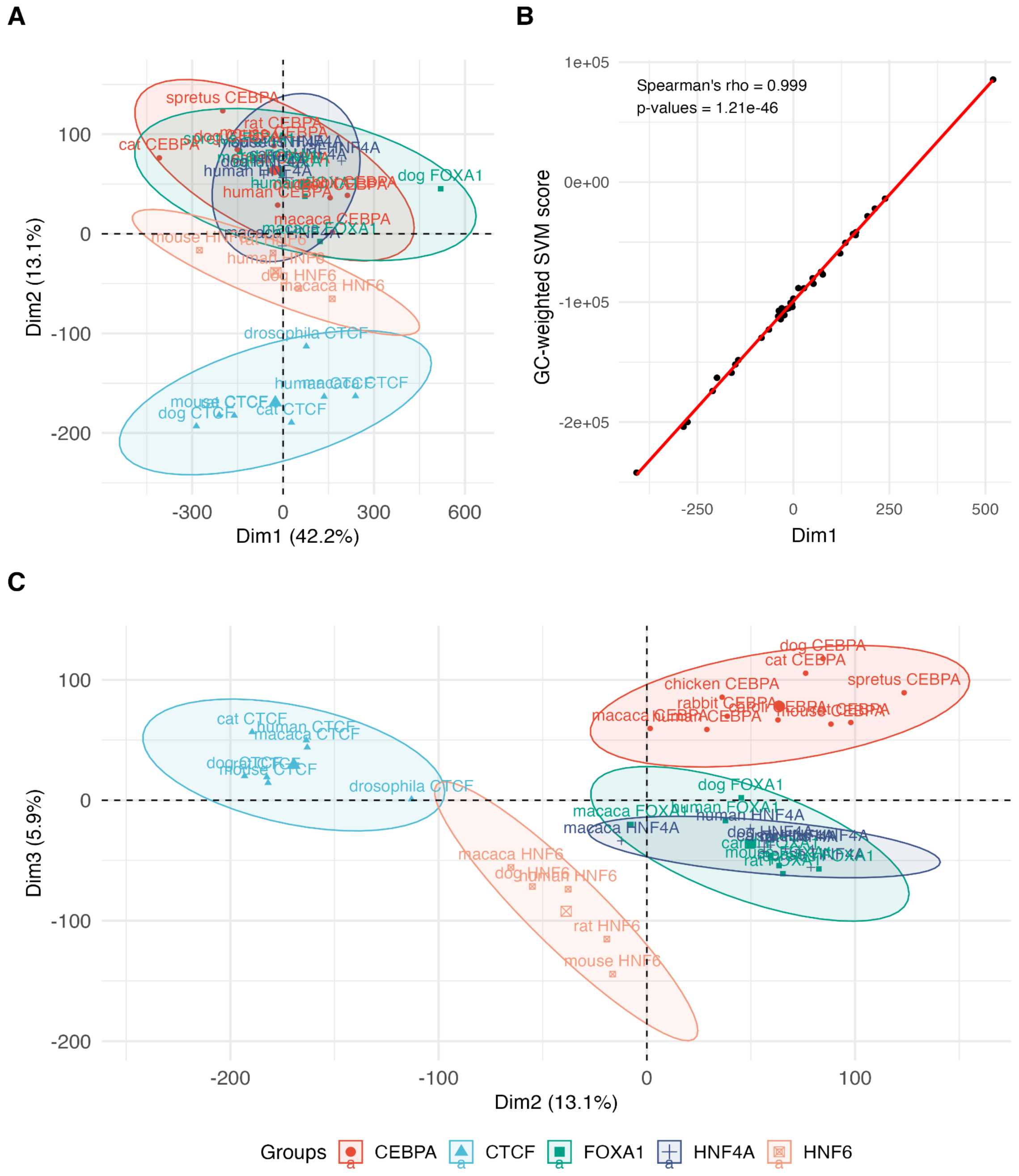


**Supplementary Figure 4: Principal component analysis of gkm-SVM models.** PCA was performed on gkm-SVM models using scores assigned to all possible 10-mers, to reflect model similarity rather than experimental replicates or sequence content. Models were trained on ChIP-seq peaks for five transcription factors across ten species (eight mammals, chicken, and *Drosophila*). Points are colored by transcription factor: CEBPA (red), CTCF (blue), FOXA1 (green), HNF4A (blue), HNF6 (orange). Ellipses represent 90% confidence intervals around each TF cluster. **A)** PCA1 versus PCA2, illustrating overall variance structure. **B)** Correlation between PCA1 and the GC-weighted SVM score, computed by multiplying the GC content of each 10-mer by its SVM score and summing across all 10-mers for each dataset. The high correlation (Spearman's rho = 0.99, p = 1.21e-46) indicates that PCA1 largely reflects variation in GC content. **C.** PCA2 versus

PCA3, highlighting TF-specific clustering and partial overlap of the co-factors FOXA1 and HNF4A.

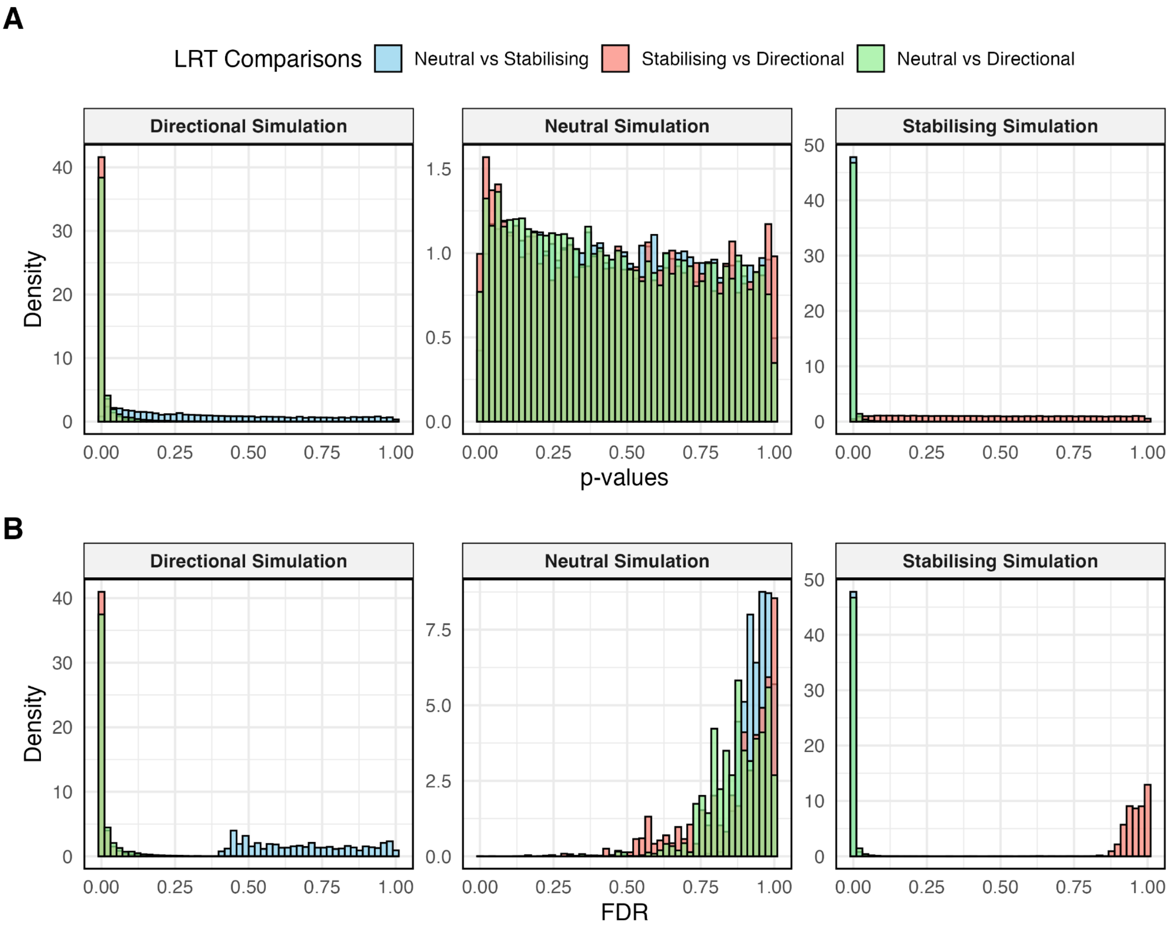


**Supplementary Figure 5: Performance of RegEvol on simulated *Drosophila* CTCF ChIP-seq peaks.**
Distributions of p-values (**A**) and FDR (**B**) from likelihood ratio tests (LRTs) are shown for three model comparisons: Neutral vs Stabilising (blue), Stabilising vs Directional (red), and Neutral vs Directional (green). Each panel contains three subpanels corresponding to sequences simulated under different evolutionary scenarios: directional selection (left; $\alpha = 25$, $\beta = 1$), neutral evolution (center; $\alpha = 1$, $\beta = 1$), and stabilising selection (right; $\alpha = 10$, $\beta = 10$). Simulations were performed on 10,000 randomly sampled Drosophila CTCF ChIP-seq peaks. Substitutions were sampled probabilistically based on their mutation probability, predicted phenotypic effect (ΔSVM), and fixation probability under the specified evolutionary model.

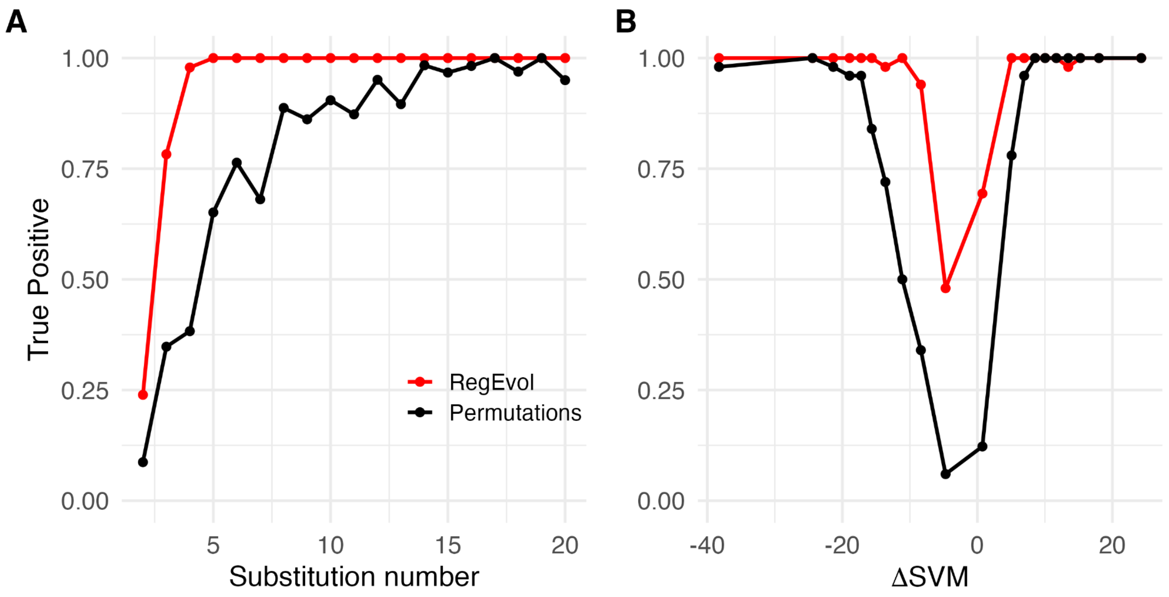


**Supplementary Figure 6: Detection performance of RegEvol under strong directional selection.**

True positive rate of peaks simulated under strong directional selection (α = 100) detected by RegEvol (red) or the Permutation Test (black) as a function of A) number of substitutions per peak and B) ΔSVM grouped by 5% quantiles. Simulations were performed on 10,000 randomly sampled Drosophila CTCF peaks. Results are shown at a false discovery rate threshold of 1%.

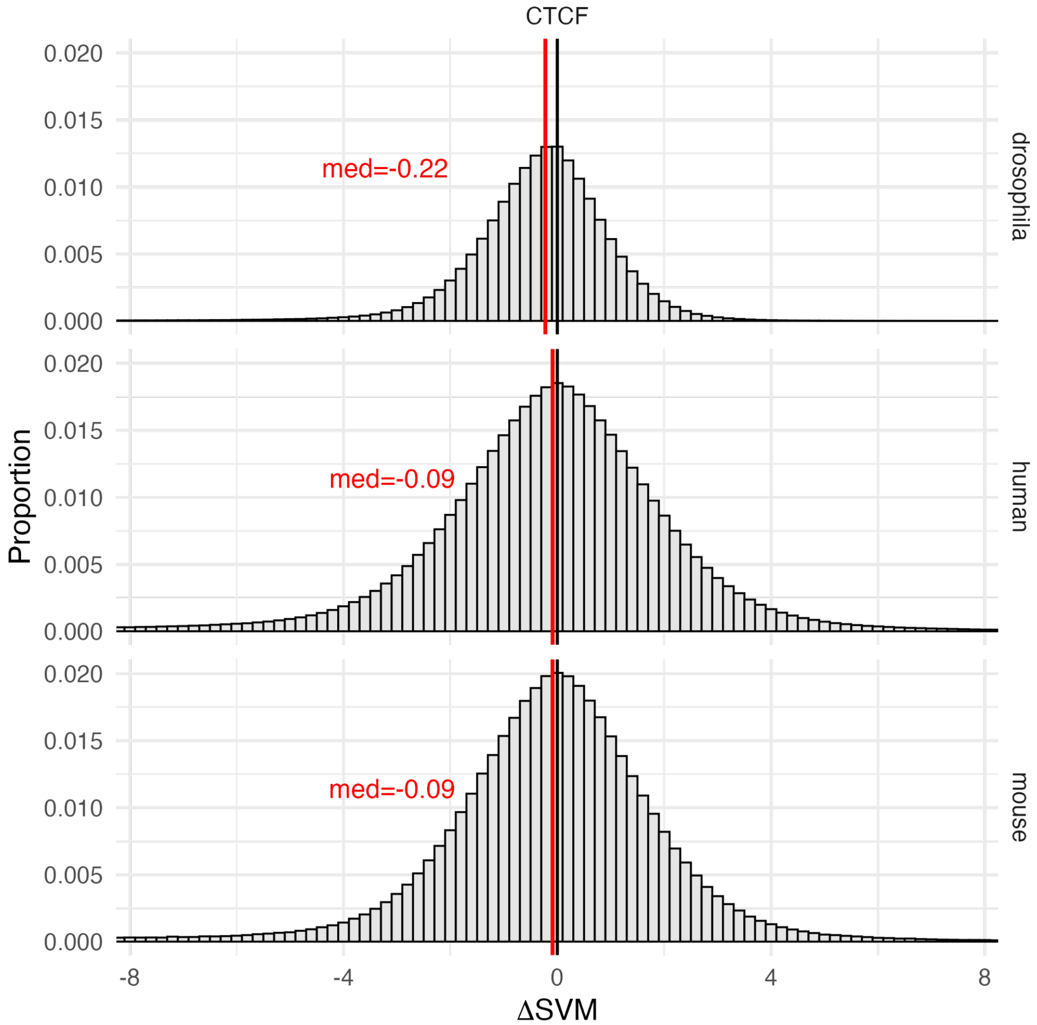


**Supplementary Figure 7: Distribution of ΔSVM from all possible point mutations in CTCF peaks.**

Distributions were obtained by introducing all possible single-nucleotide mutations in ancestral CTCF sequences. ΔSVM was computed as the change in SVM score between mutated and ancestral sequences. Panels show Drosophila (top), human (middle), and mouse (bottom). The black line indicates ΔSVM = 0, and the red line marks the slightly negative median for each species.

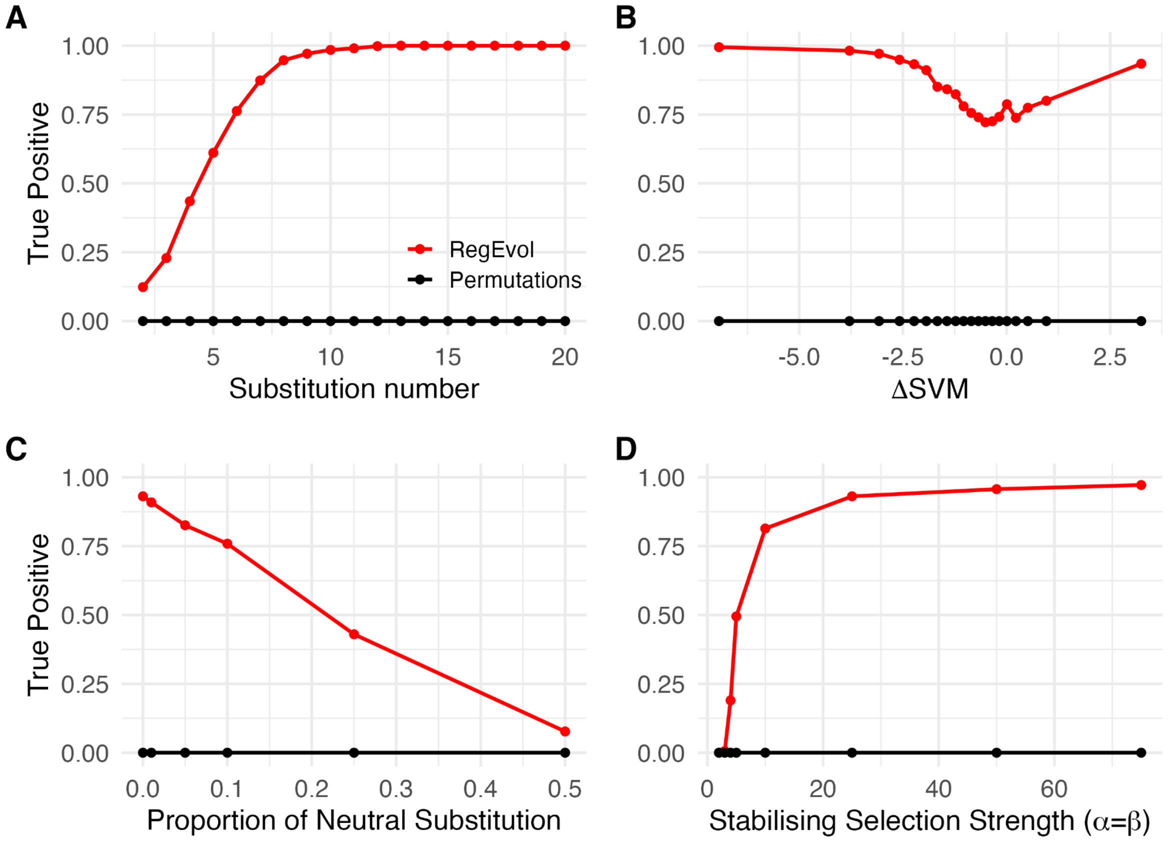


**Supplementary Figure 8: Detection performance of RegEvol under stabilising selection.**

True positive rate of peaks simulated under stabilising selection detected by RegEvol (red) or the Permutation Test (black) as a function of **A)** number of substitutions per peak, **B)** ΔSVM grouped by 5% quantiles, **C)** proportion of random substitutions, and **D)** selection strength ($\alpha = \beta$). Simulations were performed on 10,000 randomly sampled Drosophila CTCF peaks. Results are shown at a false discovery rate threshold of 1%.

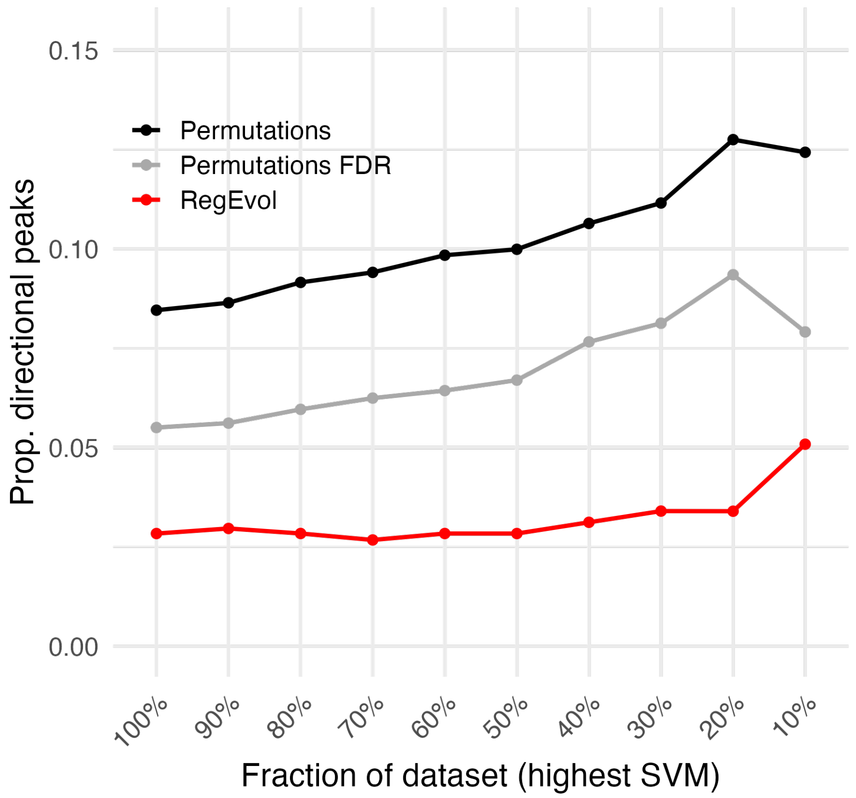


**Supplementary Figure 9: Robustness to ascertainment bias in human data.** Proportion of human liver CEBPA peaks (N = 9,958) detected under directional selection using the original Permutation Test (black lines), FDR-corrected Permutation Test (grey lines) and RegEvol (red lines) for different fractions of datasets stratified by the highest derived SVM scores. Results are shown at a false discovery rate threshold of 5%.

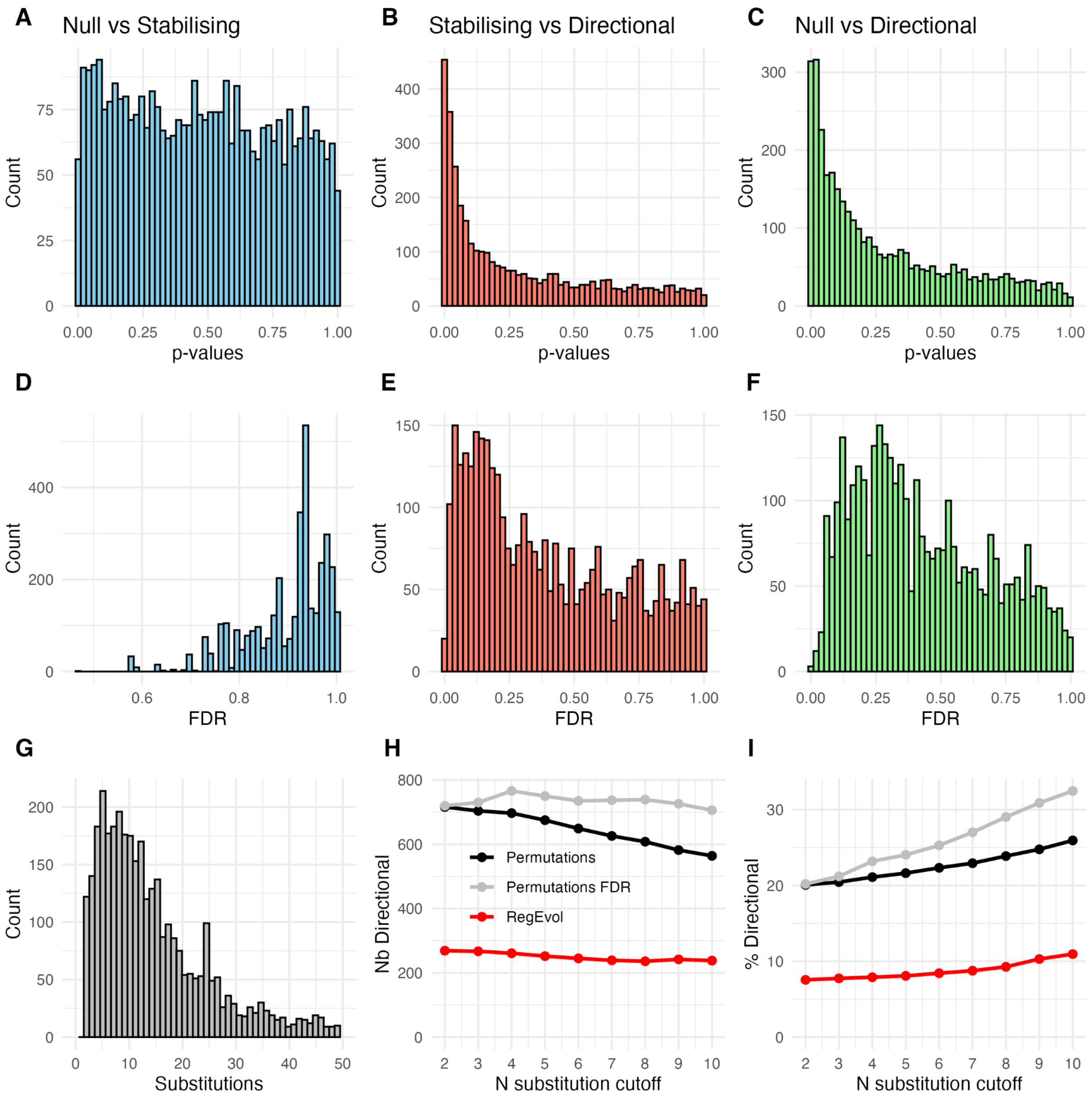


**Supplementary Figure 10: Statistical behaviour of RegEvol on *Drosophila melanogaster* CTCF ChIP-seq peaks (N = 3,566).**

**A–C)** Distributions of p-values from Likelihood Ratio Tests for different model comparisons: **A)** Null versus Stabilising, **B)** Stabilising versus Directional, **C)** Null versus Directional. **D–F)** Corresponding distributions of False Discovery Rate (FDR) for the same comparisons: **D)** Null vs Stabilising, **E)** Stabilising vs Directional, **F)** Null vs Directional. **G)** Distribution of the number of substitutions per peak. **H–I)** Effect of applying a minimum substitution cutoff followed by an FDR threshold of 5% on detection of directional selection: **H)** number of peaks detected, **I)** percentage of peaks detected by the Permutation Test (black) and RegEvol (red).

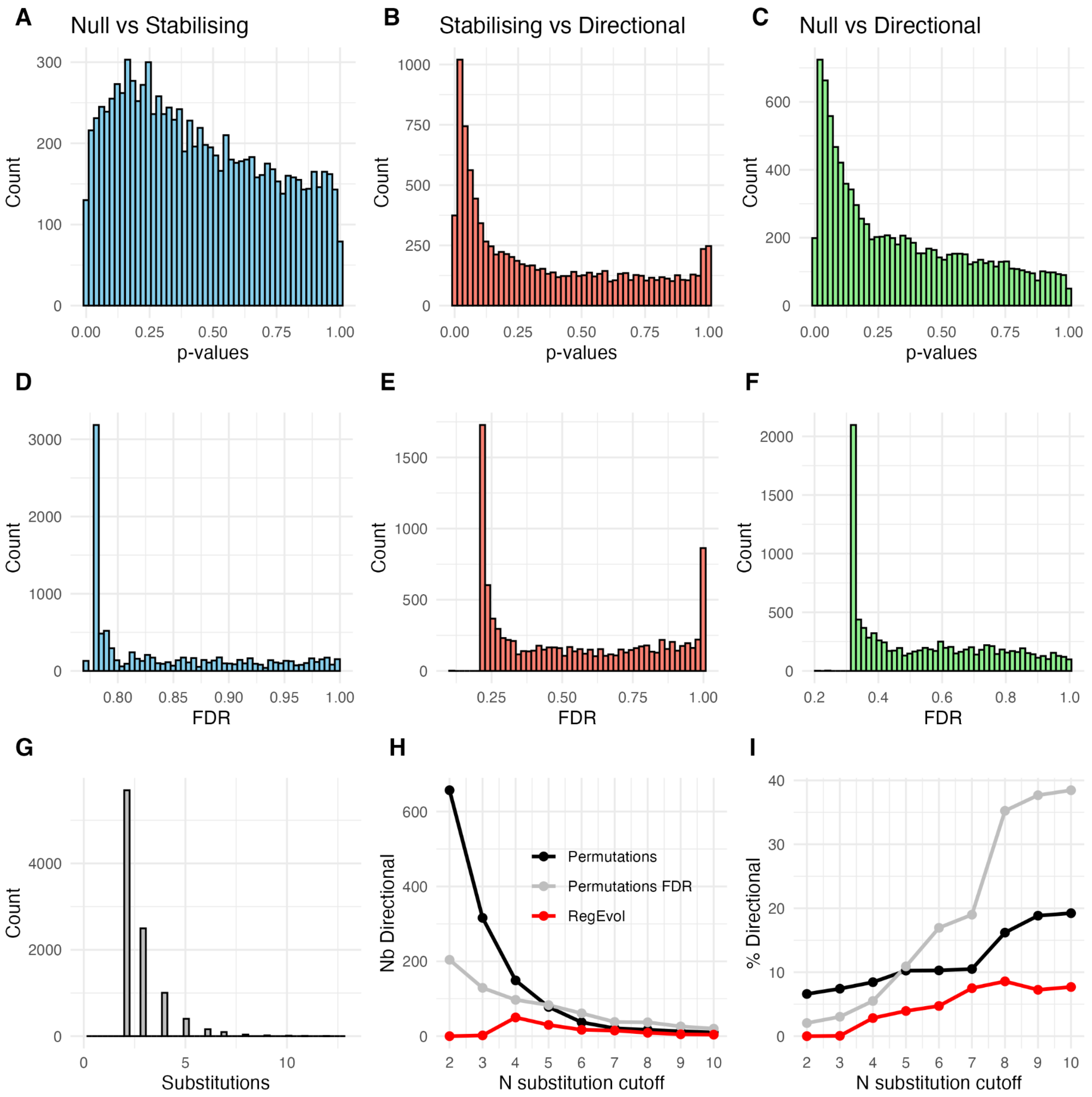


**Supplementary Figure 11: Statistical behaviour of RegEvol on *Homo sapiens* CEBPA ChIP-seq peaks (N = 9,958).**

**A–C)** Distributions of p-values from Likelihood Ratio Tests for different model comparisons: **A)** Null versus Stabilising, **B)** Stabilising versus Directional, **C)** Null versus Directional. **D–F)** Corresponding distributions of False Discovery Rate (FDR) for the same comparisons: **D)** Null vs Stabilising, **E)** Stabilising vs Directional, **F)** Null vs Directional. **G)** Distribution of the number of substitutions per peak. **H–I)** Effect of applying a minimum substitution cutoff followed by an FDR threshold of 5% on detection of directional selection: **H)** number of peaks detected, **I)** percentage of peaks detected by the Permutation Test (black) and RegEvol (red).

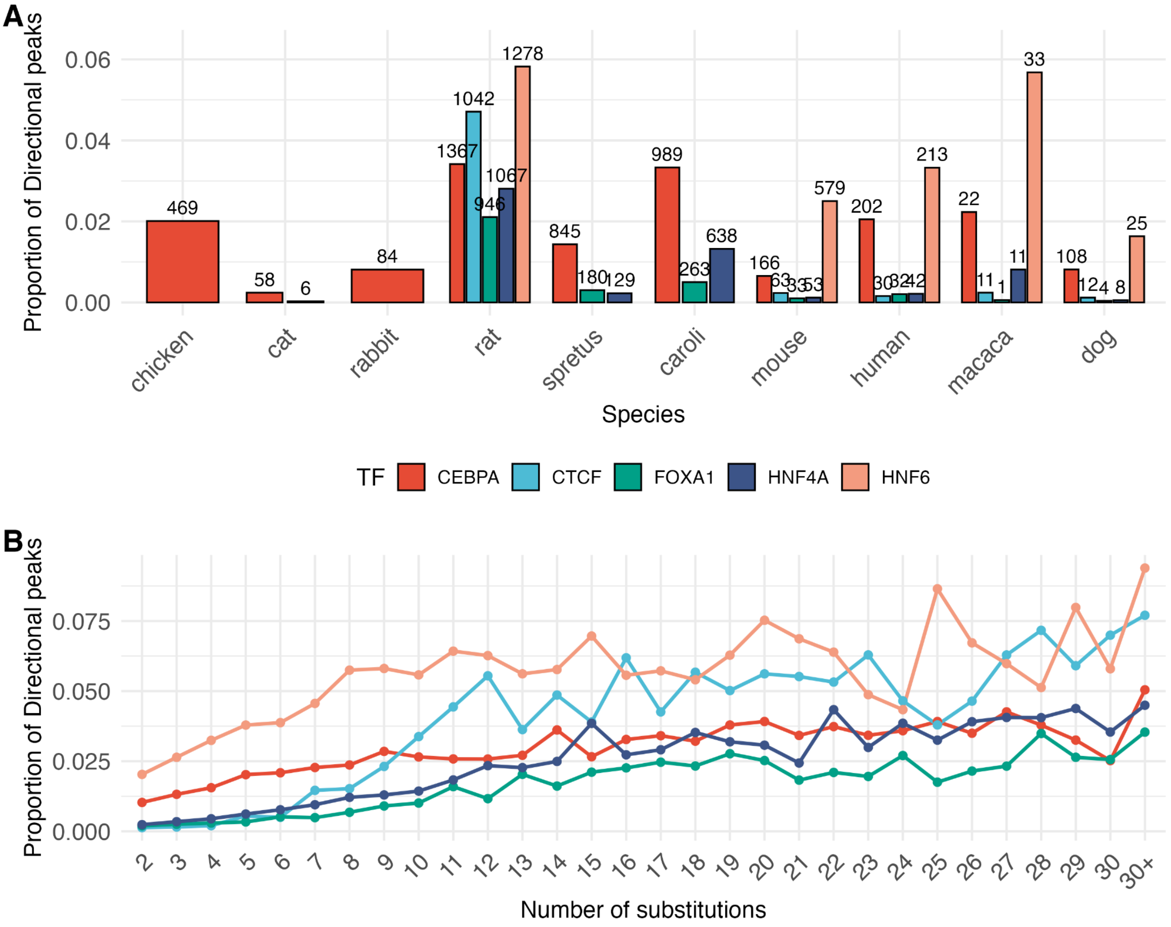


**Supplementary Figure 12: Detection of directional selection by RegEvol across vertebrate species for five transcription factors.**

**A)** Proportion of peaks inferred to be under directional selection for each dataset. The corresponding number of peaks is shown above each bar. **B)** Proportion of directional peaks for each transcription factor as a function of the number of substitutions. CEBPA (red), CTCF (blue), FOXA1 (green), HNF4A (purple), and HNF6 (orange). Only peaks with at least four substitutions are included. Results are shown at a false discovery rate threshold of 5%.

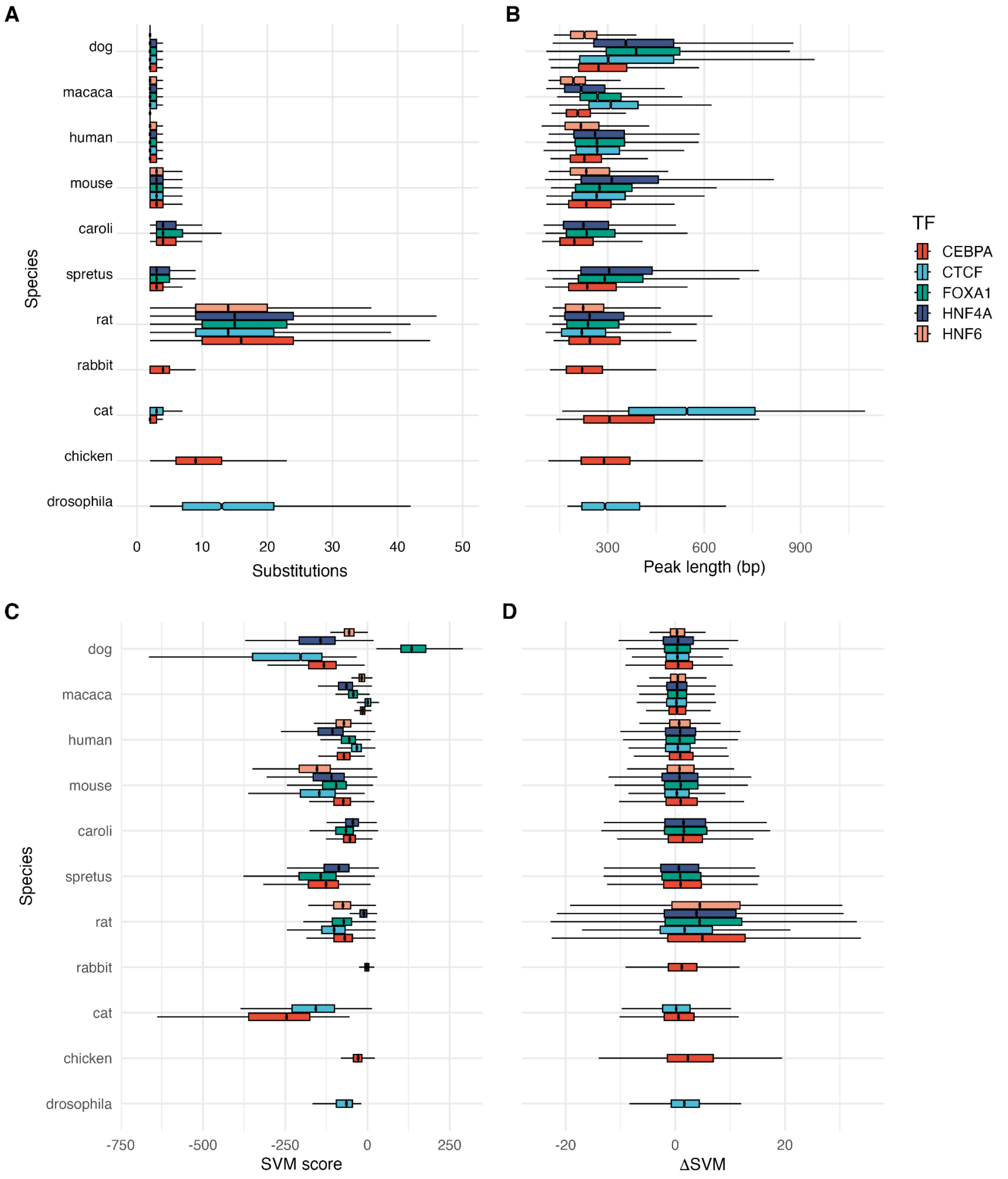


**Supplementary Figure 13: Characteristics of ChIP-seq peaks across species for five transcription factors.**

Data are shown for CEBPA (red), CTCF (blue), FOXA1 (green), HNF4A (purple), and HNF6 (orange). Boxplots summarise the following properties of peaks used in the analysis: **A)** number of substitutions per peak, **B)** peak length in base pairs, **C)** SVM score of the focal sequences, **D)** ΔSVM between focal and ancestral sequences.

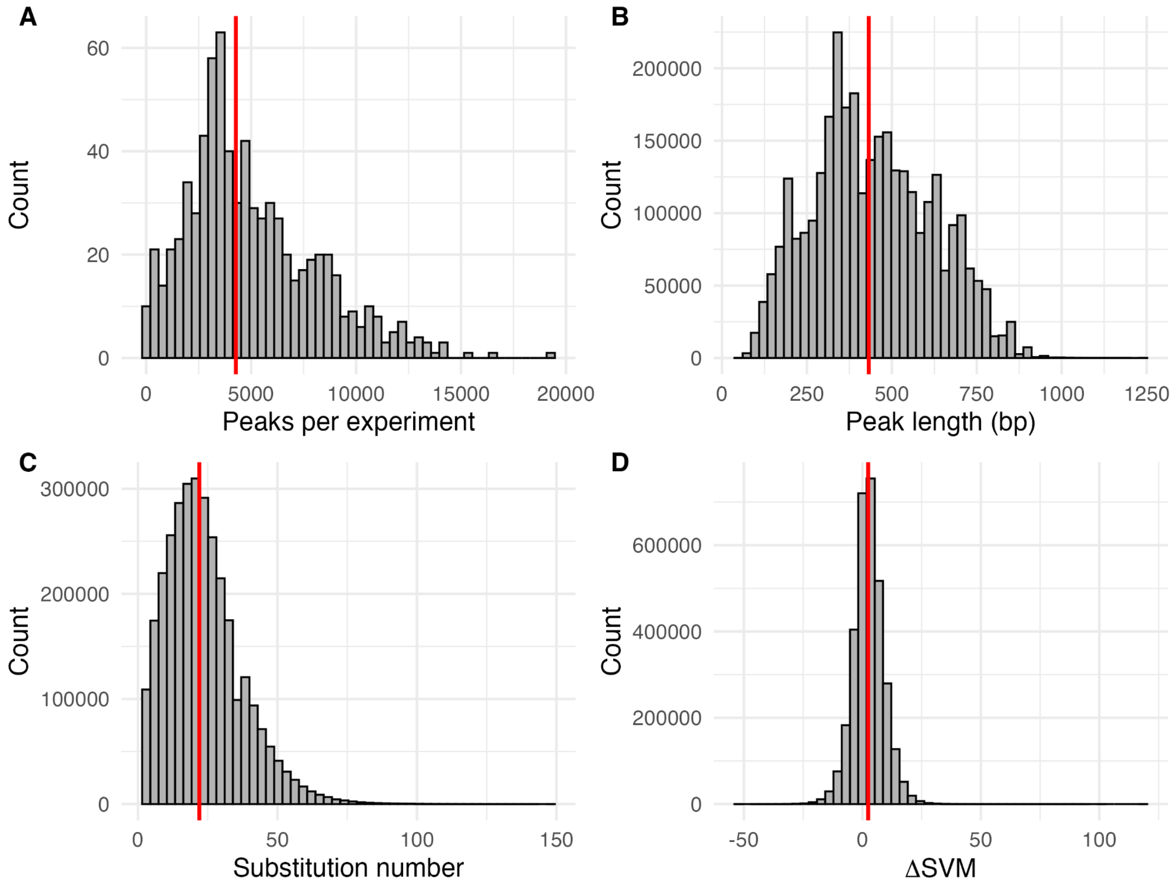


**Supplementary Figure 14: Characteristics of Drosophila melanogaster modERN peaks.**

Distributions of peak properties across all 740 experiments, including all peaks (N = 3,190,372), are shown for: **A)** number of peaks per experiment, **B)** peak length in base pairs, **C)** number of substitutions per peak, **D)** ΔSVM between focal and ancestral sequences. Vertical red lines indicate the median for each distribution.

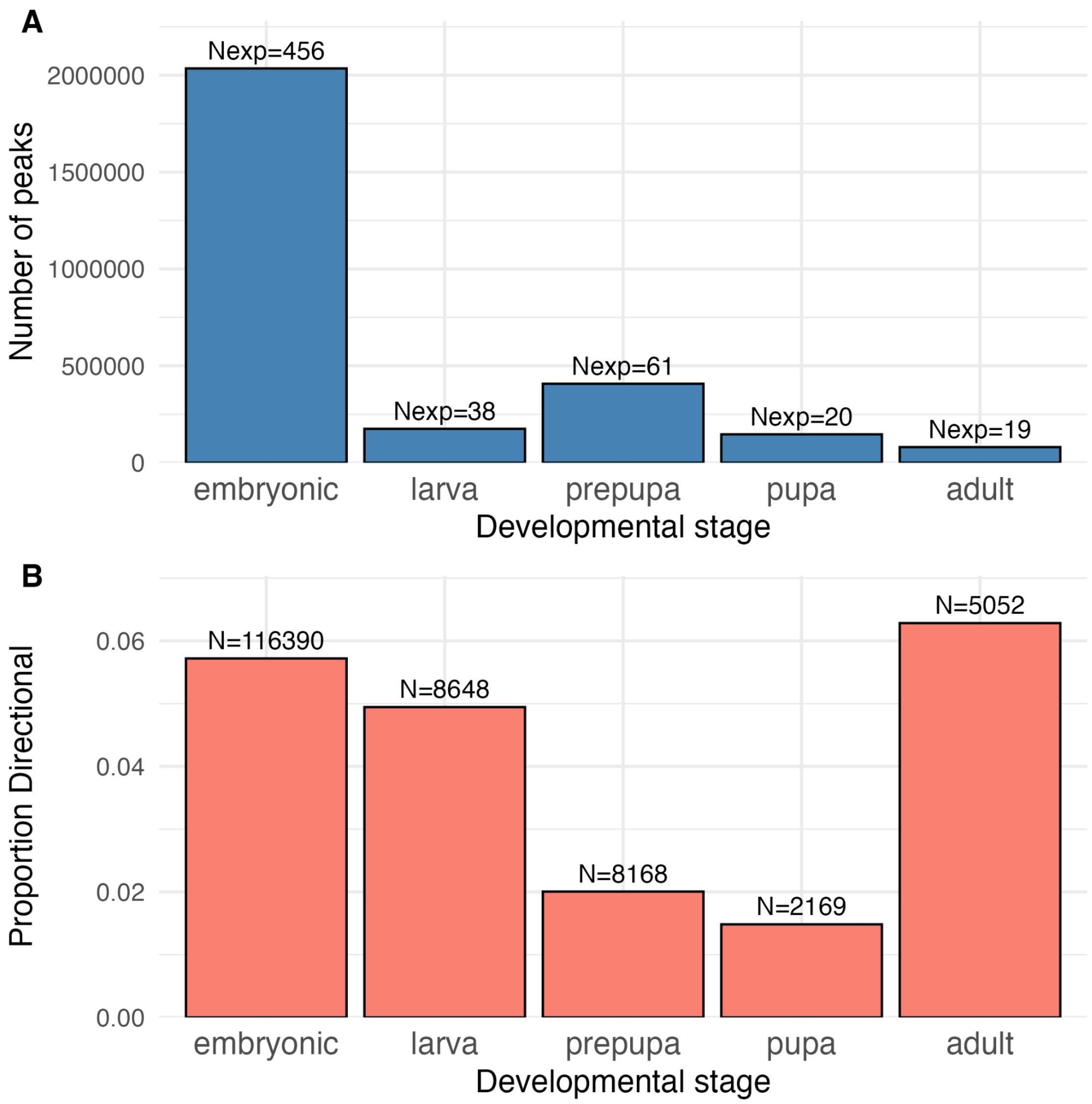


**Supplementary Figure 15: *Drosophila melanogaster* modERN peaks per developmental stage.**

**A)** Number of analysed peaks for each developmental stage. Developmental stages were directly extracted from modENCODE annotations. The number above each bar indicates the number of experiments contributing to that stage. **B)** Proportion of peaks inferred to be under directional selection by RegEvol for each developmental stage. The number above each bar indicates the corresponding number of peaks. Results are shown at a false discovery rate threshold of 5%.

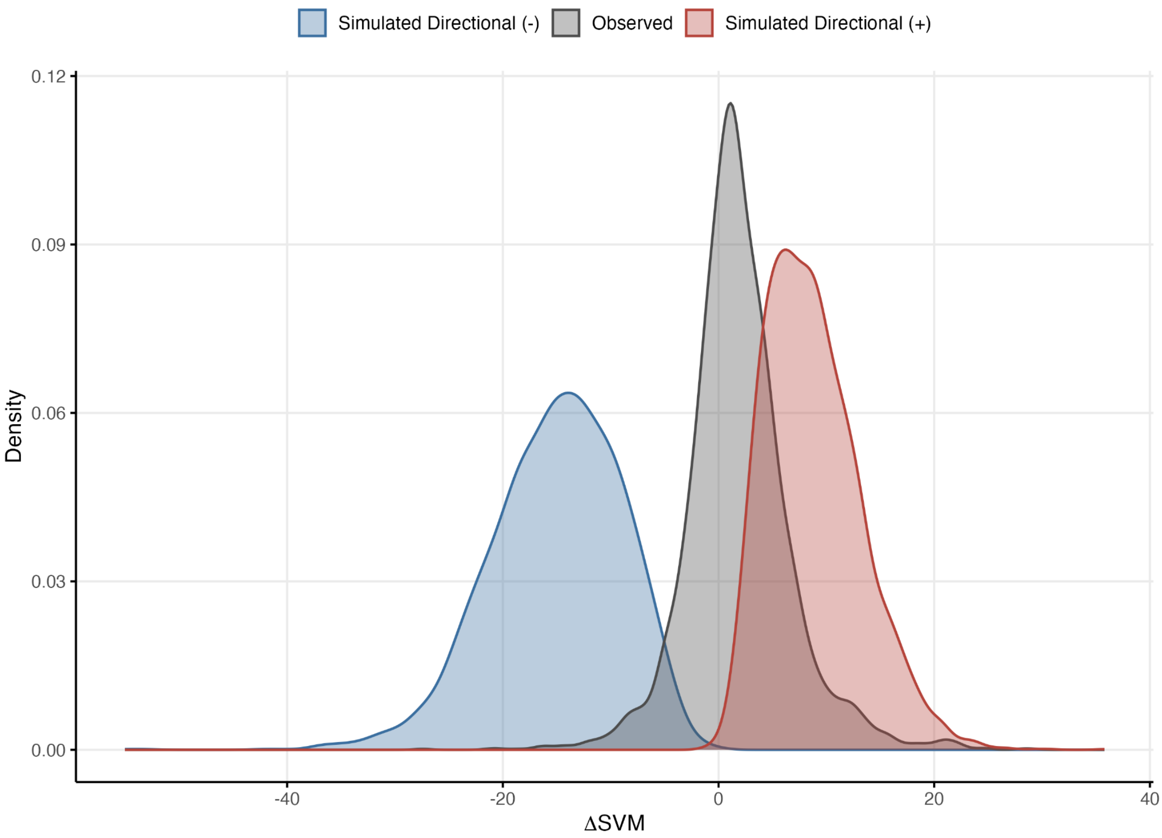


**Supplementary Figure 16: Detection limits of negative directional selection.**
Ancestral sequences of *Drosophila melanogaster* CTCF peaks were evolved by simulation under directional selection towards decreased binding affinity (blue) and towards increased binding affinity (red). The resulting ΔSVM distributions are compared to the distribution of ΔSVM values actually observed within *D. melanogaster* CTCF peaks (grey). Only the upper tail of the simulated decreasing directional distribution (blue) overlaps with the observed range of ΔSVM (overlap coefficient = 0.16), whereas the simulated increasing directional distribution (red) overlaps a larger range of observed values (overlap coefficient = 0.50).

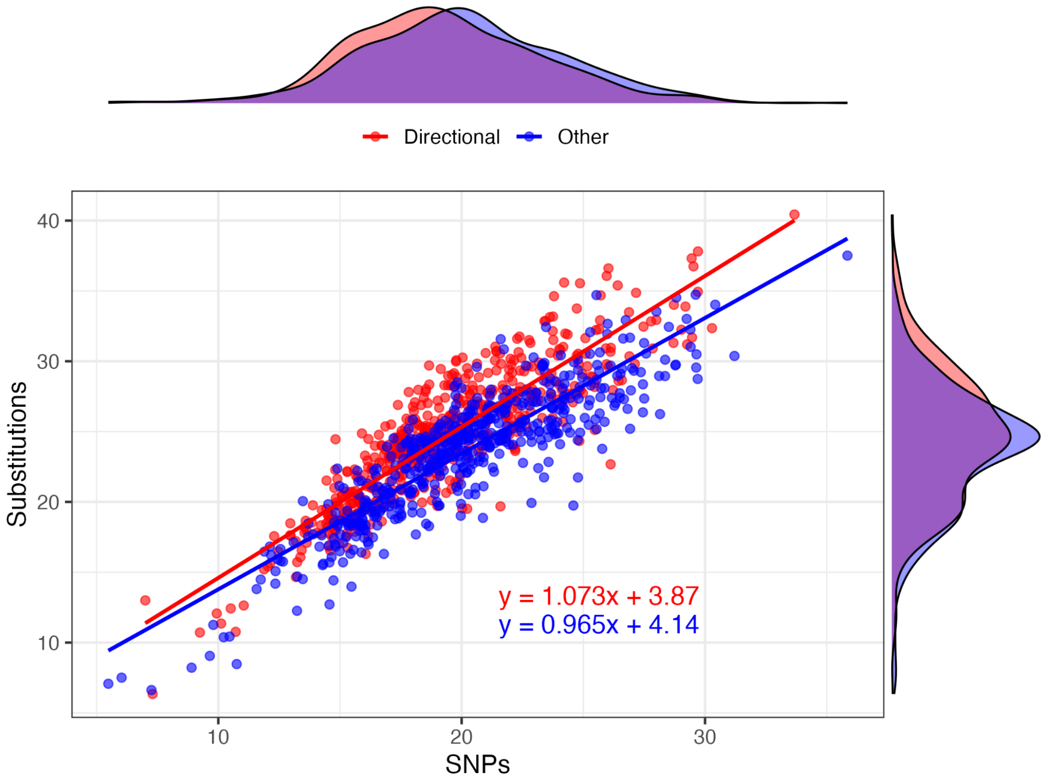


**Supplementary Figure 17: *Drosophila melanogaster* substitutions vs SNPs.** Fixed substitutions were determined from the alignment between *D. melanogaster* and *D. simulans*, and polymorphisms were obtained from the Drosophila Genetic Reference Panel (DGRP). Only positions overlapping *D. melanogaster* modERN peaks were considered (Materials & Methods). Mean values were then calculated for each experiment for peaks detected under directional selection (red) and for all other peaks (blue). Directional peaks have significantly more substitutions (mean = 24.67 vs 23.54; Wilcoxon test, $p = 1.8 \times 10^{-4}$) and fewer SNPs (mean = 19.38 vs 20.11; Wilcoxon test, $p = 2.9 \times 10^{-4}$) compared to other peaks.

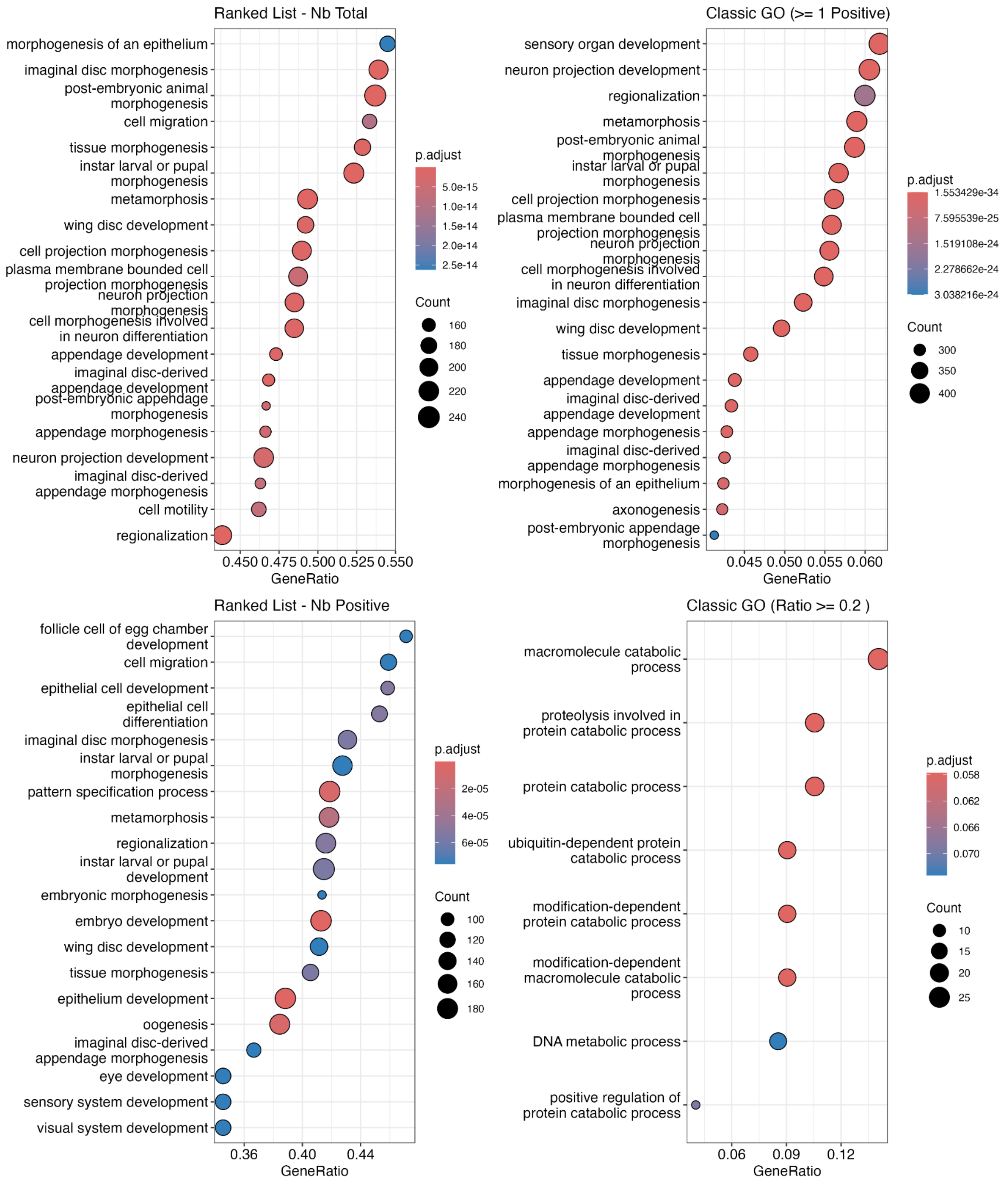


**Supplementary Figure 18: Gene Ontology Enrichment.**

Gene Ontology (GO) terms enriched among genes associated with *Drosophila melanogaster* modERN peaks. Associations between peaks and genes were obtained from the modERN consortium based on genomic proximity (Materials & Methods). Only the first 20 enriched GO terms are shown. **A)** Genes ranked by total number of associated peaks. **B)** Genes with at least one directional peak compared with all genes with at least one associated peak. **C)** Genes ranked by the total number of directional peaks. **D)** Genes ranked by the ratio of directional to total peaks (restricted to genes with ≥10 associated peaks). Point colour indicates FDR-adjusted p-values and point size reflects the number of genes associated with each GO term.

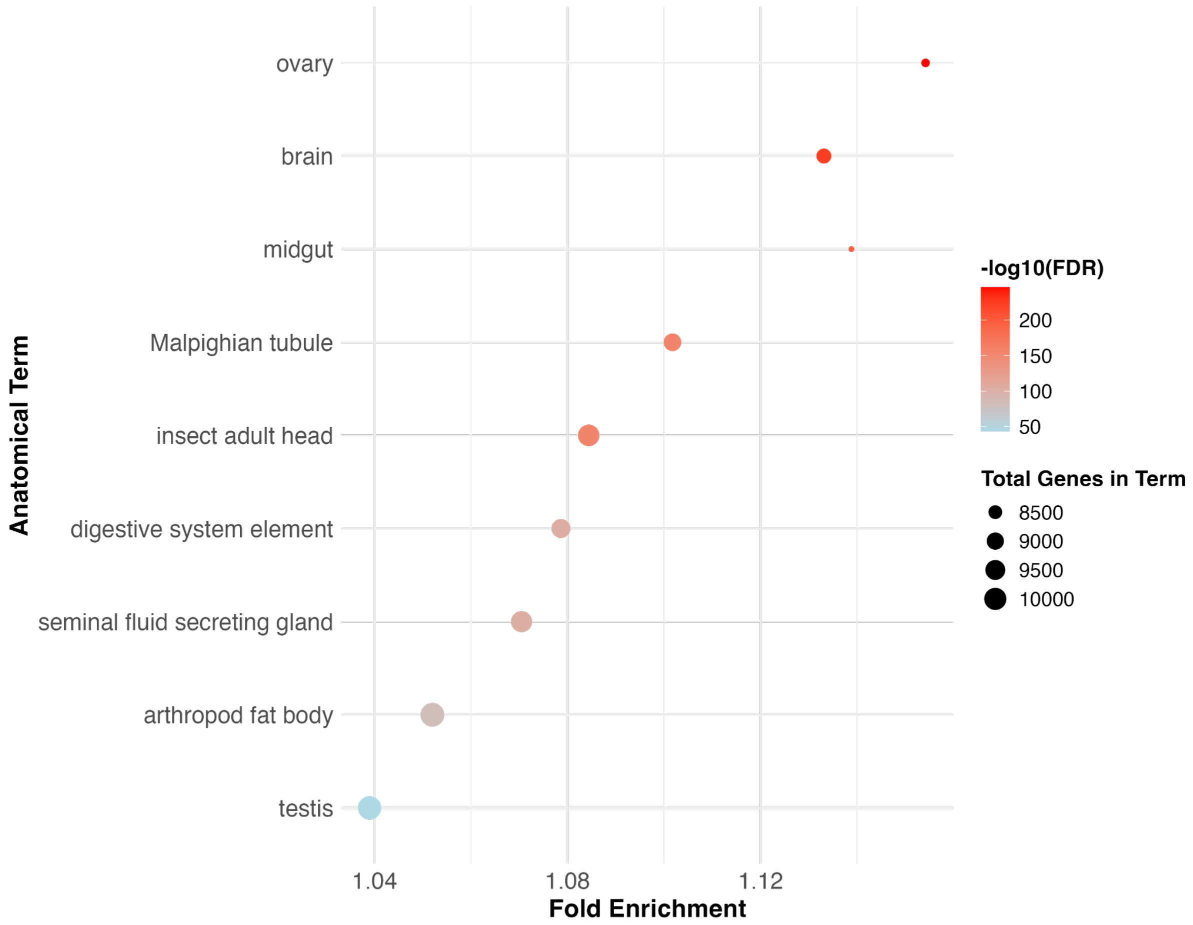


**Supplementary Figure 19: TopAnat Drosophila**

Anatomical terms enriched among genes with a directional-to-total peak ratio ≥0.2 compared with all genes with associated peaks (restricted to genes with ≥10 associated peaks). TopAnat identifies anatomical structures where a gene set is preferentially expressed using integrated RNA-seq expression data (Materials & Methods). Only terms with FDR-adjusted p-values ≤0.05 are shown. Point size reflects the number of genes associated with each anatomical term.

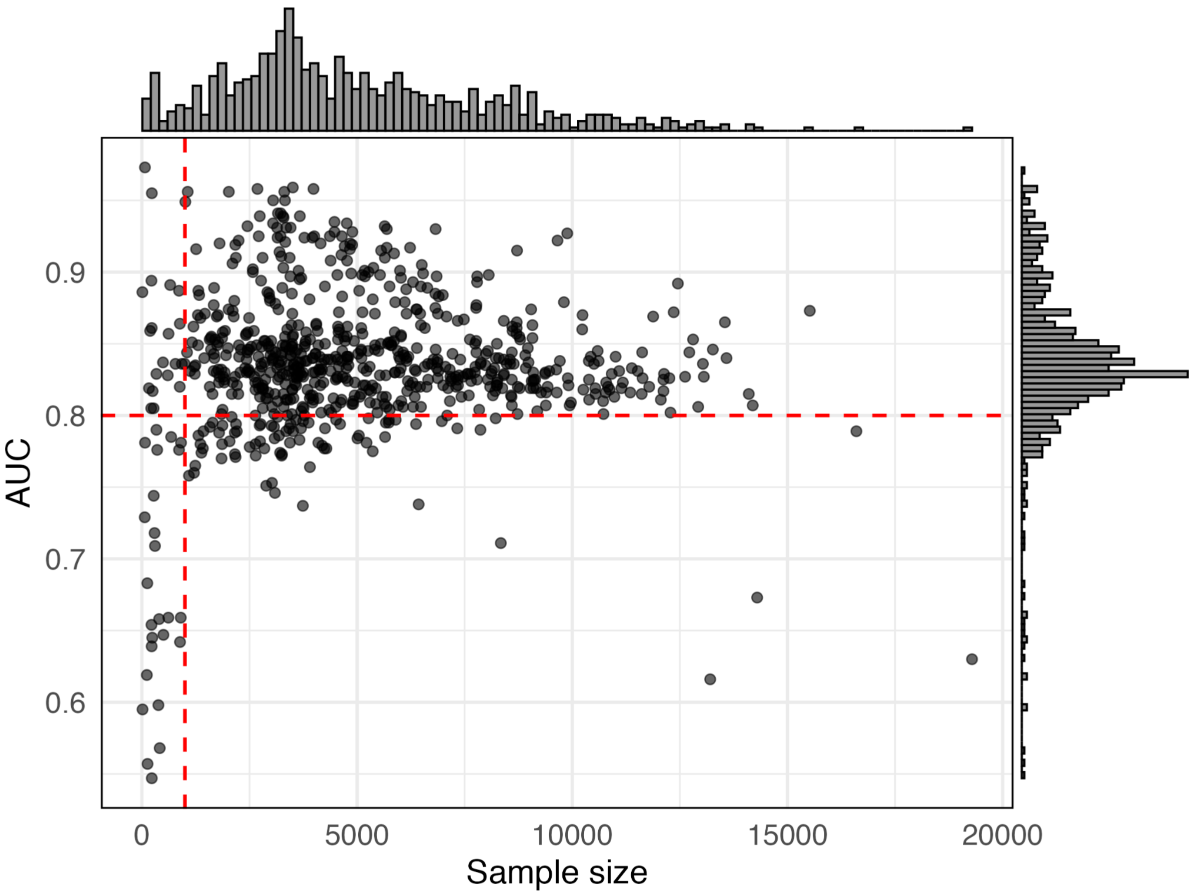


**Supplementary Figure 20: Model performance of Drosophila modERN experiments.** Area under the curve (AUC) values were calculated from 5-fold cross-validation of gkm-SVM models for each experiment. The AUC measures the ability of the model to distinguish TF-bound regulatory sequences from random genomic sequences, with 1 indicating perfect discrimination and 0.5 representing random performance. Sample size represents the number of peaks reported by the modERN consortium. Experiments with fewer than 1,000 peaks or with an AUC below 0.8 (dotted red lines) were excluded from downstream analyses, resulting in 615 experiments out of 740 (Materials & Methods).

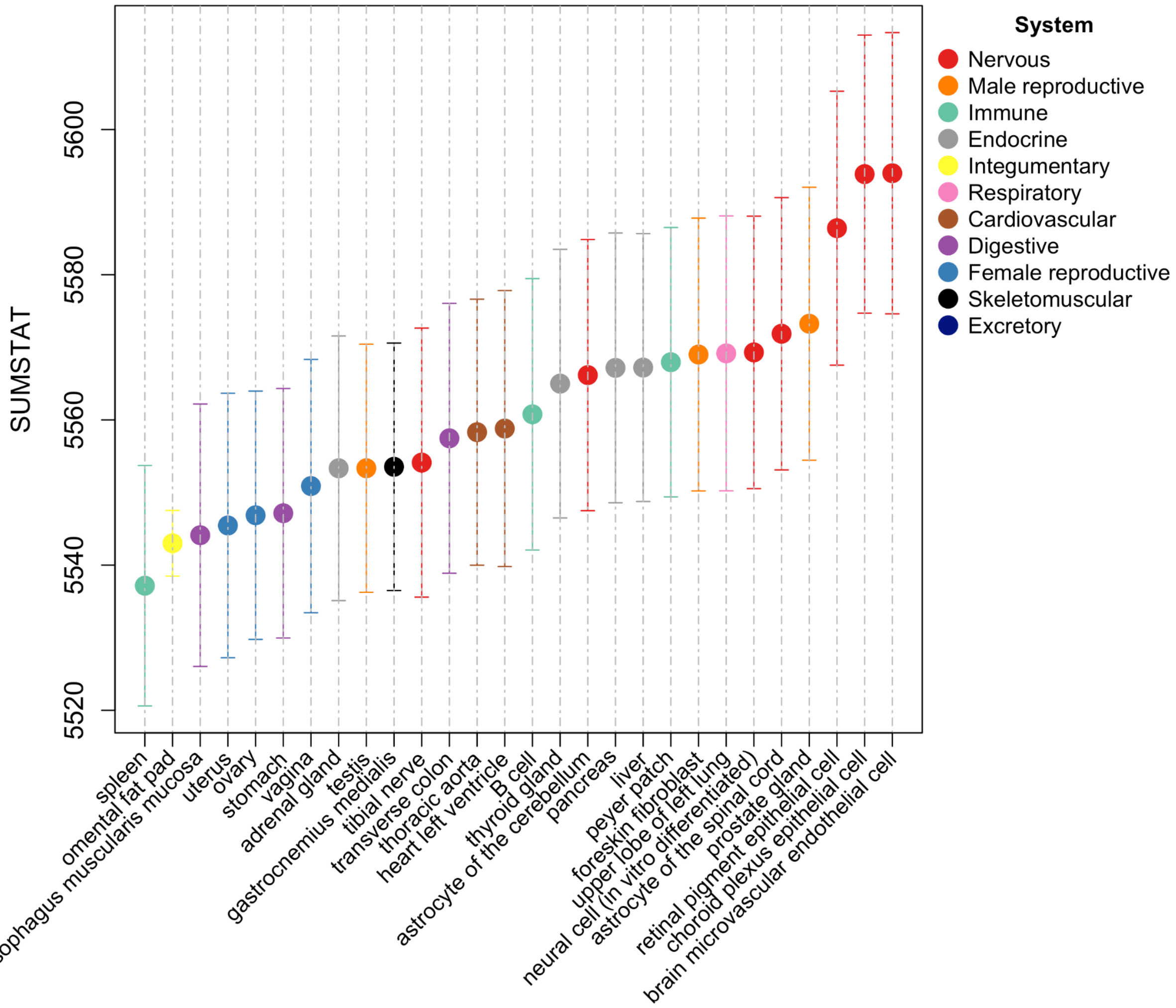


**Supplementary Figure 21: Tissue-level aggregation of directional selection on chimpanzee homologs of human CTCF peaks.**

Median SUMSTAT scores across 10,000 resamplings of 5,000 CTCF peaks per tissue. Bars indicate the standard deviation across resamplings. SUMSTAT scores were calculated following the framework from Daub et al. (2017) by taking the fourth-root of the per-peak Δlog-likelihood (Directional minus Neutral model) and summing across all peaks within each tissue. Tissues are sorted by median SUMSTAT and colored according to organ system.
